# Growth-dependent activation of protein kinases suggests a mechanism for measuring cell growth

**DOI:** 10.1101/610469

**Authors:** Akshi Jasani, Tiffany Huynh, Douglas R. Kellogg

## Abstract

In all cells, progression through the cell cycle occurs only when sufficient growth has occurred. Thus, cells must translate growth into a proportional signal that can be used to measure and transmit information about growth. Previous genetic studies in budding yeast suggested that related kinases called Gin4 and Hsl1 could play roles in mechanisms that measure bud growth; however, interpretation of the data was complicated by the use of gene deletions that cause complex terminal phenotypes. Here, we used the first conditional alleles of Gin4 and Hsl1 to more precisely define their functions. We show that excessive bud growth during a prolonged mitotic delay is an immediate consequence of inactivating Gin4 and Hsl1. Thus, acute loss of Gin4 and Hsl1 causes cells to behave as though they cannot detect that bud growth has occurred. We further show that Gin4 and Hsl1 undergo gradual hyperphosphorylation during bud growth that is dependent upon growth and correlated with the extent of growth. Moreover, gradual hyperphosphorylation of Gin4 during bud growth requires binding to anionic phospholipids that are delivered to the growing bud. While alternative models are possible, the data suggest that signaling lipids delivered to the growing bud generate a growth-dependent signal that could be used to measure bud growth.

## Introduction

Key cell cycle transitions occur only when sufficient growth has occurred. To enforce this dependency relationship, cells must convert growth into a proportional signal that triggers cell cycle progression when it reaches a threshold. However, the molecular mechanisms by which cells generate proportional signals to measure and limit cell growth have remained deeply mysterious.

In budding yeast, growth occurs in 3 distinct intervals that are characterized by different rates and patterns of growth (McCusker *et al.* 2007; Goranov *et al.* 2009; Ferrezuelo *et al.* 2012; Leitao and Kellogg 2017). The first interval occurs during G1 phase and is characterized by slow growth over the entire cell surface. The second interval is initiated at the end of G1 phase when a new daughter cell emerges and undergoes polar growth. The third interval is initiated at entry into mitosis and is marked by a switch from polar bud growth to growth that occurs more widely over the bud surface. Bud growth then continues throughout mitosis. The distinct size and shape of a yeast cell is ultimately defined by the extent of growth during each of these intervals.

Several observations suggest that mechanisms that control the duration and extent of bud growth play a major role in control of cell size (Leitao and Kellogg 2017). First, little growth occurs during G1 phase. For example, cells growing in rich nutrients increase in size by only 20% during G1 phase. Rather, most growth occurs during bud growth in mitosis, and the rate of growth in mitosis is approximately 3-fold faster than growth during the other intervals. Therefore, failure to tightly control the duration and extent of bud growth, particularly during mitosis, would have large consequences for cell size.

Additional evidence that bud growth is tightly controlled comes from analysis of the effects of nutrient availability on cell growth and size. Growth of cells in a poor nutrient source causes a large reduction in average cell size (Johnston *et al.* 1977; 1979). However, growth in poor nutrients has little effect on cell size at completion of G1 phase (Bean *et al.* 2006; Leitao and Kellogg 2017). Furthermore, mutant cells that lack all known regulators of cell size in G1 phase show robust nutrient modulation of cell size (Jorgensen and Tyers 2004a). In contrast, poor nutrients cause a large decrease in the extent of growth in mitosis, which causes daughter cells to complete cytokinesis at a substantially reduced size (Hartwell and Unger 1977; Leitao and Kellogg 2017). Furthermore, the duration of growth in mitosis is increased in poor nutrients, which suggests that cells compensate for slow bud growth by increasing the duration of growth (Leitao and Kellogg 2017). These observations point to the existence of mechanisms that measure and modulate both the duration and extent of bud growth to ensure that mitotic progression is linked to cell growth.

A model in which the extent of bud growth is tightly controlled requires a molecular mechanism for measuring growth. Here, we searched for proteins that could play a role in measuring bud growth during mitosis. A candidate-based approach identified 3 related kinases: Gin4, Hsl1 and Kcc4, which we refer to as Gin4-related kinases. Similar kinases are found in all eukaryotes. Genetic analysis reaching back over 30 years has suggested that Gin4-related kinases could play roles in control of cell growth and size (Young and Fantes 1987; Ma *et al.* 1996; Altman and Kellogg 1997; Okuzaki *et al.* 1997; Barral *et al.* 1999). For example, loss of Gin4-related kinases in both budding yeast and fission yeast causes excessive growth during a prolonged delay in mitotic progression (Young and Fantes 1987; Ma *et al.* 1996; Okuzaki *et al.* 1997; Barral *et al.* 1999). A model that could explain this phenotype is that Gin4-related kinases play roles in measuring bud growth. In this case, cells that lack Gin4-related kinases would behave as though they cannot detect that bud growth has occurred. However, an alternative model is that aberrant growth caused by loss of Gin4-related kinases is an indirect consequence of mitotic delays. For example, there is evidence that loss of Gin4-related kinases causes defects in positioning the mitotic spindle, which could cause chromosome segregation defects and associated mitotic delays that lead to abnormally prolonged growth (Fraschini *et al.* 2006; Grava *et al.* 2006; Gihana *et al.* 2018). In this case, growth defects would be a secondary consequence of mitotic spindle defects.

A major problem with distinguishing models is that previous genetic analysis of Gin4-related kinases utilized gene deletions that cause severe terminal phenotypes. This has made it difficult to discern which aspects of the phenotype are an immediate and direct consequence of loss of function, versus aspects of the phenotype that are the result of secondary defects accumulated over multiple generations. An additional major limitation of previous studies is that they provided limited information on the physiological signals that control Gin4-related kinases. If Gin4-related kinases are involved in measuring growth, then their activity must in some way be mechanistically linked to growth. Previous studies found that Gin4 is gradually hyperphosphorylated and activated during mitosis (Altman and Kellogg 1997). Since bud growth occurs gradually throughout mitosis, this observation suggests that Gin4 activity could be a readout of growth; however, the data are correlative and activation of Gin4 or Hsl1 has not been analyzed in the context of cell growth.

Here, we created the first conditional alleles of Gin4-related kinases and used them to show that defects in control of bud growth are an immediate and direct consequence of inactivating Gin4-related kinases. We further show that Gin4-related kinases influence the duration and extent of growth during metaphase, and that they do so partly via regulation of Cdk1 inhibitory phosphorylation. Finally, we show that Gin4 related kinases undergo gradual phosphorylation during bud growth that is dependent upon bud growth and correlated with the extent of bud growth. While alternative models remain possible, the data suggest a model in which Gin4-related kinases generate and/or relay growth-dependent signals that could be used to measure the extent of bud growth.

## Results

### Gin4-related kinases are required for normal control of bud growth during mitosis

Gin4 and Hsl1 are the most important Gin4-related kinases in budding yeast. Loss of either kinase alone causes defects in control of bud growth, whereas loss of both causes severe defects (Barral *et al.* 1999; Longtine *et al.* 2000). Loss of the Gin4 paralog Kcc4 has little effect. We therefore focused on Gin4 and Hsl1. To avoid the complications associated with long term inactivation of the Gin4-related kinases, we created auxin-inducible degron (AID) versions of Gin4 and Hsl1, which allowed us to define the immediate effects of inactivation (Nishimura *et al.* 2009). A strain carrying AID-tagged versions of both *GIN4* and *HSL1* had no size defects in the absence of auxin (**Fig. S1 A**). Addition of auxin before mitosis in synchronized cells caused a large reduction in levels of Gin4-AID protein within 30 minutes (**Fig. S1 B**), as well as a delay in mitotic progression (**Fig. S1 C**).

We first tested how conditional inactivation of Gin4 and Hsl1 influences bud growth and mitotic progression via live analysis of single cells. We included a fluorescently tagged spindle pole protein to monitor the duration of metaphase and anaphase (see Materials and Methods and (Leitao and Kellogg 2017)). The spindle poles in wild type and *gin4-AID hsl1-AID* cells were marked with different fluorescent tags, which allowed simultaneous imaging of both strains under identical conditions.

We analyzed the effects of *gin4-AID* or *hsl1-AID* alone, as well as the combined effects of *gin4-AID hsl1-AID*. Cells were released from a G1 arrest and auxin was added immediately before initiation of bud emergence, which ensured that Gin4 and Hsl1 were depleted by the time of mitotic entry. Bud size and mitotic spindle dynamics were then analyzed at 3-minute intervals to determine how loss of Gin4 and Hsl1 influenced bud growth and the duration of mitosis. Examples of wild type and *gin4-AID hsl1-AID* cells are shown in **Fig. 1 and Video 1**. Both cells initiated bud emergence at nearly the same time, but the wild type cell completed bud growth and exited mitosis while the *gin4-AID hsl1-AID* cell remained delayed in metaphase as the bud continued to grow. The *gin4-AID hsl1-AID* cell eventually completed mitosis, but at a substantially larger bud size than the wild type control cell. The daughter bud was more elongated in the *gin4-AID hsl1-AID* cell, which indicated a defect in control of polar growth.

**Figure 1:**
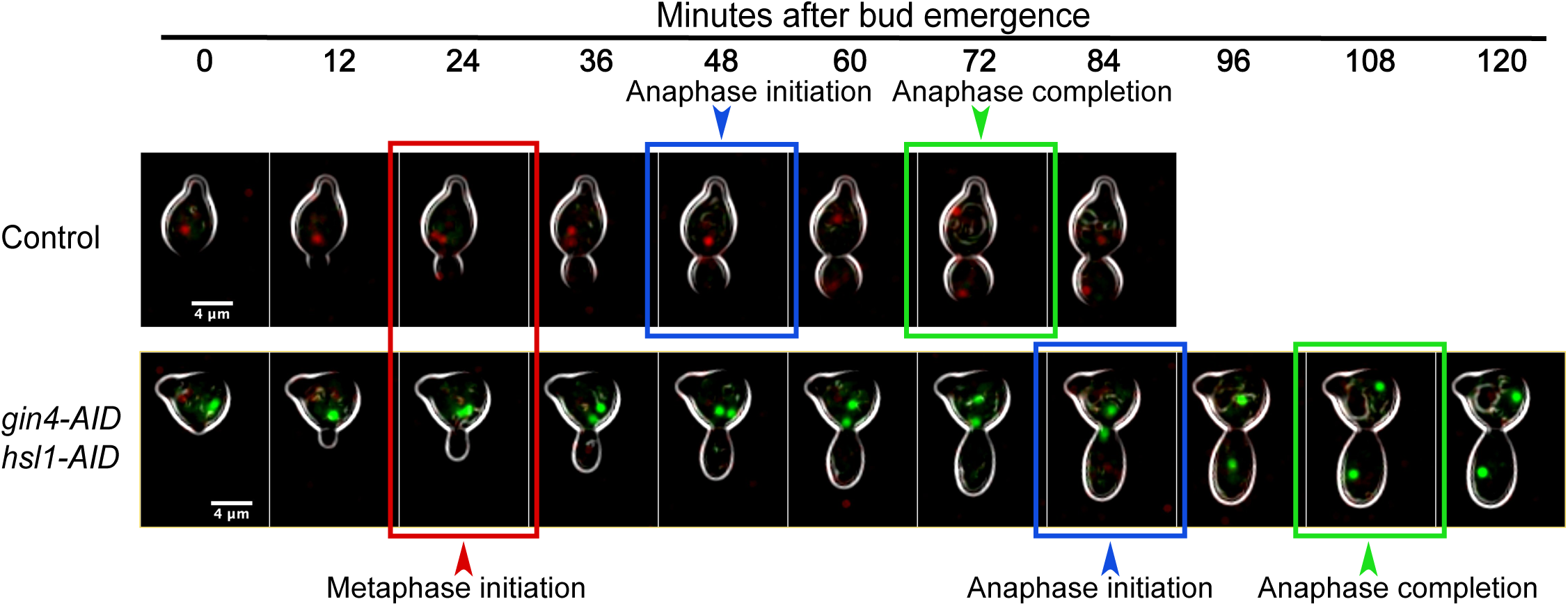
Gin4 and Hsl1 are required for normal control of cell growth and size in mitosis. Control cells and *gin4-AID hsl1-AID* cells were differentially marked with fluorescently-tagged mitotic spindle poles. Thus, control cells express *SPC42-mRuby2*. while the *gin4-AID hsl1-AID* cells express *SPC42-GFP*. Both strains include 2 copies of the *TIR1* gene. Cells growing in CSM were arrested with α factor and then mixed together before releasing from the arrest. Auxin was added to 0.5 mM at 20 minutes after release from arrest, which corresponds to approximately 30 minutes before bud emergence. Cells were then imaged at 3-minute intervals by confocal microscopy at 27°C. Bud emergence was used to set the zero timepoint. Key mitotic transitions are highlighted for each strain.

Quantitative analysis of multiple cells showed that destruction of Gin4 and/or Hsl1 caused an increase in the duration of metaphase but had little effect on the duration of anaphase (**Fig. 2 A**). Destruction of Gin4 and Hsl1 also caused an increase in bud size at completion of each mitotic interval (**Fig. 2 B**). *gin4-AID hsl1-AID* did not cause significant effects on the growth rate of the daughter cell (**Fig. S2 A**). The effects of *gin4-AID* and *hsl1-AID* were not additive (**Fig. 2 A and B**), which was surprising because *hsl1*Δ and *gin4*Δ have strong additive effects on cell size and shape (**Fig. S2 B** and (Barral *et al.* 1999)). This issue is addressed below. Inactivation of *gin4-AID* caused polar bud growth to continue in mitosis, whereas inactivation of *hsl1-AID* did not. This effect was quantified by measuring axial ratios of daughter buds at completion of anaphase (**Fig. 2 C**).

**Figure 2:**
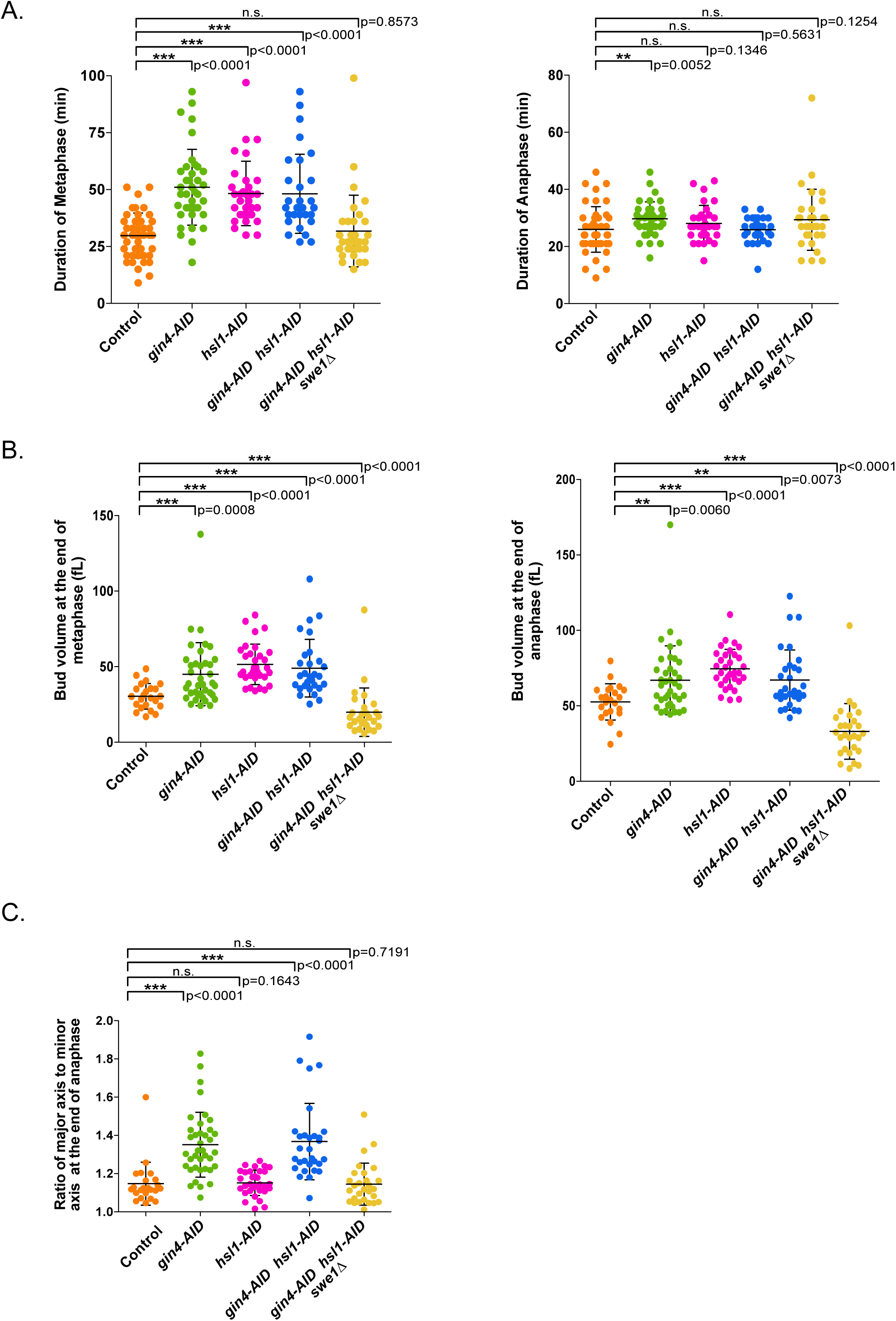
Gin4 and Hsl1 are required for normal control of cell growth and size in mitosis. Cells of the indicated genotypes were released from a G1 arrest and analyzed by confocal microscopy as described for Fig. 1. All strains included 2 copies of the *TIR1* gene. **(A)** Scatter plots showing the duration of metaphase and anaphase. **(B)** Scatter plots showing bud size at completion of metaphase and anaphase. **(C)** Scatter plots showing the ratio of major axis to minor axis of the bud at completion of anaphase. For panels **A, B, C**, the mean and standard deviation for each strain are shown.

Previous studies found that *gin4*Δ and *hsl1*Δ cause defects in cytokinesis that lead to formation of clumps of interconnected cells (Ma *et al.* 1996; Okuzaki *et al.* 1997; Barral *et al.* 1999). Consistent with this, we observed that *gin4-AID hsl1-AID* caused a failure in cell separation in nearly all cells at the end of the first cell cycle following the addition of auxin (**Video 2**). Previous studies also found that *gin4*Δ and *hsl1*Δ cause defects in spindle positioning (Fraschini *et al.* 2006; Grava *et al.* 2006; Gihana *et al.* 2018). We observed only a few defects in spindle positioning during the first cell division after addition of auxin to *gin4-AID hsl1-AID* cells. However, in the second cell division many cells showed aberrant movement of the metaphase spindle into the daughter cell before anaphase (**Video 2**). Thus, defects in spindle orientation appear to be a secondary consequence of inactivating Gin4 and Hsl1.

### Gin4-related kinases are required for normal control of mother cell growth

In wild type cells, little growth of the mother cell occurs after bud emergence (McCusker *et al.* 2007; Ferrezuelo *et al.* 2012). In *gin4-AID hsl1-AID* cells, however, we found that mother cell growth often continued throughout the interval of daughter cell growth. Example plots of mother and daughter cell size as a function of time for wild type and *gin4-AID hsl1-AID* cells are shown in **Fig. 3 A**. Quantitative analysis revealed that *gin4-AID* and *hsl1-AID* had additive effects upon mother cell growth so that most *gin4-AID hsl1-AID* cells underwent abnormal mother cell growth (**Fig. 3 B**).

**Figure 3:**
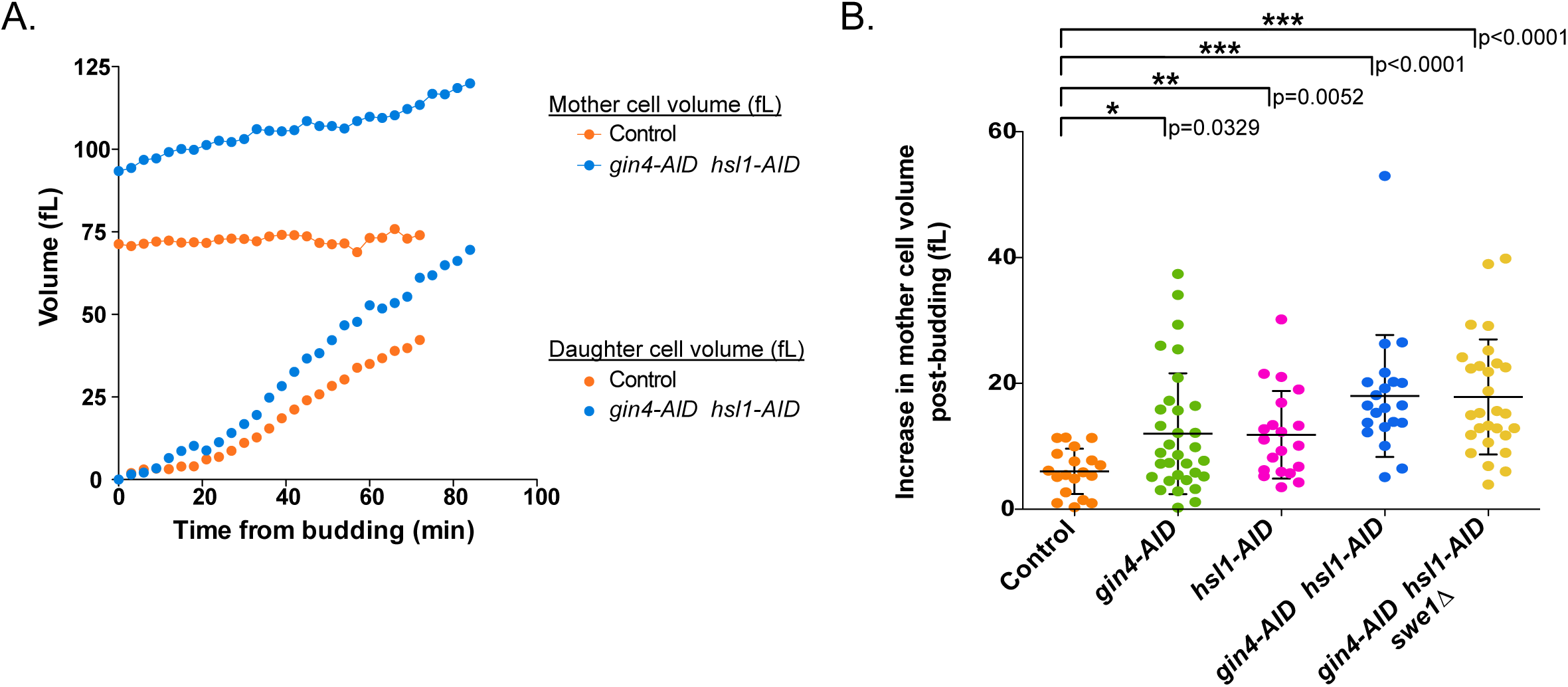
Gin4 and Hsl1 are required for normal control of mother cell growth. Cells of the indicated genotypes were released from a G1 arrest and analyzed by confocal microscopy as described for Fig. 1. **(A)** A representative plot of mother and daughter cell size as a function of time. In each case, the daughter cell is the daughter of the mother cell shown in the same plot. **(B)** Scatter plots showing the net increase in mother cell volume from the time of bud emergence to completion of anaphase. The plot shows the mean and standard deviation for each strain.

Previous work has shown that daughter cell size is correlated with mother cell size (Schmoller *et al.* 2015; Leitao and Kellogg 2017). Thus, large mother cells give birth to large daughter cells. As a result, the increased size of mothers in *gin4-AID hsl1-AID* cells would be expected to drive increased daughter cell size in subsequent cell divisions, which could lead to gradually increasing defects in cell growth and size. Therefore, the role of Gin4 and Hsl1 in control of mother cell growth could help explain why prolonged loss of Gin4 and Hsl1 causes strong additive effects on cell growth and size. We found that the effects caused by *gin4-AID hsl1-AID* increased substantially with prolonged incubation in the presence of auxin, consistent with a model in which the terminal phenotype caused by *gin4*Δ *hsl1*Δ is partially the result of defects that accumulate over multiple cell cycles (**Fig. S3**).

### Gin4-related kinases influence the duration of growth during metaphase via inhibitory phosphorylation of Cdk1

Genetic analysis has shown that Gin4-related kinases are negative regulators of the Wee1 kinase, which phosphorylates and inhibits mitotic Cdk1 (Ma *et al.* 1996; Longtine *et al.* 2000). The budding yeast homolog of Wee1 is referred to as Swe1. Additional studies have shown that Swe1 influences the timing of mitotic entry as well as the duration of metaphase by regulating this inhibitory phosphorylation of Cdk1 (Harvey and Kellogg 2003; Lianga *et al.* 2013; Leitao *et al.* 2019). We therefore tested whether Gin4 and Hsl1 influence the duration of metaphase and cell size via Swe1. To do this, we analyzed the effects of *swe1*Δ on bud growth and mitotic duration in *gin4-AID hsl1-AID* cells. This revealed that *swe1*Δ eliminated the prolonged metaphase delay caused by loss of Gin4 and Hsl1 (**Fig. 2 A**). Furthermore, *swe1*Δ caused daughter buds in *gin4-AID hsl1-AID* cells to complete metaphase and anaphase at sizes smaller than the wild type control cells (**Fig. 2 B**), and it eliminated the bud elongation caused by *gin4-AID* (**Fig. 2 C**). As reported previously, *swe1*Δ caused reduced growth rate, which is thought to be due to the decreased size of mother cells (**Fig. S2 A**) (Leitao *et al.* 2019).

Several observations demonstrated that Gin4 and Hsl1 do not work solely via Swe1. Previous studies found that *swe1*Δ cells have a shorter metaphase than wild type cells (Lianga *et al.* 2013; Leitao *et al.* 2019). Here, we found that *swe1*Δ reduced the duration of metaphase in *gin4-AID hsl1-AID* cells, but it did not make the duration of metaphase in these cells shorter than metaphase in wild type cells, indicating that acute loss of Gin4 and Hsl1 still influenced the duration of metaphase in *swe1*Δ cells. In addition, *swe1*Δ did not fully rescue growth defects caused by *gin4*Δ *hsl1*Δ (**Fig. S2 B**). Finally, *swe1*Δ did not eliminate inappropriate growth of mother cells in *gin4-AID hsl1-AID* cells (**Fig. 3 B**). Together, these observations demonstrate that Gin4 and Hsl1 have Swe1-dependent and Swe1-independent functions.

### Gin4 and Hsl1 are required for full hyperphosphorylation of Swe1

We next investigated how Gin4 and Hsl1 control Swe1. In previous work, we showed that Swe1 undergoes complex regulation in mitosis (Sreenivasan and Kellogg 1999; Harvey *et al.* 2005; 2011). In early mitosis, Cdk1 phosphorylates Swe1 on Cdk1 consensus sites, which activates Swe1 to bind and inhibit Cdk1. This form of Swe1, which we refer to as partially hyperphosphorylated Swe1, works in a systems-level mechanism that maintains a low level of Cdk1 during metaphase. Further phosphorylation events drive full hyperphosphorylation of Swe1, leading to release of Cdk1 and inactivation of Swe1. Swe1 is proteolytically destroyed at the end of mitosis; however, mutants that block Swe1 destruction have no effect on mitotic progression, so the function of Swe1 destruction remains unknown (Thornton and Toczyski 2003; Raspelli *et al.* 2011).

A previous study suggested that *hsl1*Δ causes defects in phosphorylation of Swe1 but did not provide sufficient resolution of differently phosphorylated forms of Swe1 to determine which events were affected (Shulewitz *et al.* 1999). Here, we found that conditional inactivation of *gin4-AID* and *hsl1-AID* before mitosis caused a failure in full hyperphosphorylation of Swe1 (**Fig. 4**). Similarly, addition of auxin to asynchronous *gin4-AID hsl1-AID* cells caused loss of fully hyperphosphorylated Swe1 within 30 minutes (**Fig. S4**). These data show that Gin4 and Hsl1 are required for generation of the fully hyperphosphorylated inactive form of Swe1, consistent with genetic data showing that Gin4-related kinases are negative regulators of Swe1.

**Figure 4:**
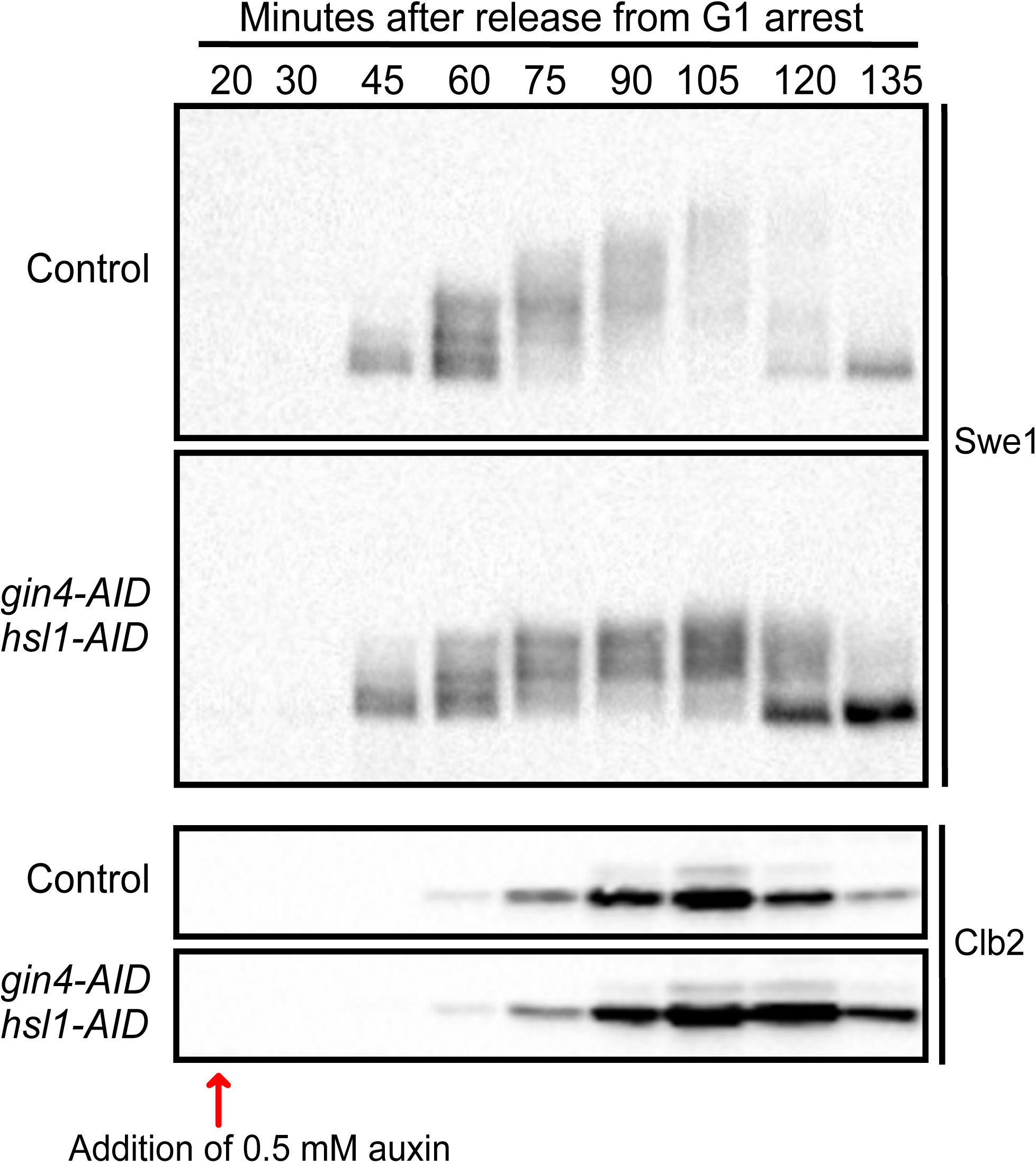
Gin4 and Hsl1 are required for full hyperphosphorylation of Swe1. Control cells and *gin4-AID hsl1-AID* cells growing in YPD were released from a G1 arrest at 25°C and 0.5 mM auxin was added to both strains 20 min after release. Both strains included 2 copies of the *TIR1* gene. Samples were taken at the indicated intervals and the behavior of Swe1 and Clb2 was analyzed by western blot.

### Hyperphosphorylation of Gin4 and Hsl1 is correlated with the extent of bud growth

We previously found that Gin4 undergoes gradual hyperphosphorylation and activation during bud growth, reaching peak hyperphosphorylation and peak kinase activity in late mitosis as growth ends (Altman and Kellogg 1997). Thus, gradual phosphorylation and activation of Gin4 seemed to correlate with gradual bud growth, which suggests that Gin4 could be activated by growth-dependent signals that provide a molecular readout of the extent of bud growth. To begin to test this hypothesis, we first carried out additional experiments to determine whether hyperphosphorylation of Gin4 and Hsl1 is correlated with bud growth. To do this, we took advantage of the fact that the durations of both metaphase and anaphase are increased when cells are growing slowly on a poor carbon source, even as the extent of growth in mitosis is reduced (Leitao and Kellogg 2017; Leitao *et al.* 2019). In other words, cells in poor carbon spend more time growing in both metaphase and anaphase, but complete mitosis at a smaller daughter cell size. Therefore, we reasoned that if hyperphosphorylation of Gin4 and Hsl1 correlates with the extent of bud growth, the duration of hyperphosphorylation should be increased in cells growing in poor carbon, while the extent of hyperphosphorylation should be reduced. Alternatively, if phosphorylation of Gin4 and Hsl1 is more closely linked to a mitotic event unrelated to growth, such as positioning of the mitotic spindle, one might expect that the timing and/or extent of hyperphosphorylation would not be influenced by carbon source.

Wildtype cells growing in rich carbon (2% glucose) or poor carbon (2% glycerol, 2% ethanol) were released from a G1 arrest and phosphorylation of Gin4 and Hsl1 was assayed by western blot to detect phosphorylation events that cause electrophoretic mobility shifts. The same samples were also probed for the mitotic cyclin Clb2 as a marker for mitotic duration. Cells growing in poor carbon showed delayed mitotic entry and a prolonged mitosis compared to cells growing in rich carbon, as previously described (**Fig. 5 A, B**) (Leitao and Kellogg 2017; Leitao *et al.* 2019). Gin4 is present throughout the cell cycle, while Hsl1 is synthesized anew before mitosis and destroyed at the end of mitosis (Altman and Kellogg 1997; Burton and Solomon 2000). In both carbon sources, hyperphosphorylation of Gin4 and Hsl1 increased gradually during mitosis, with peak phosphorylation occurring near peak Clb2 levels. In addition, the interval during which hyperphosphorylation occurred was prolonged in poor carbon. Previous studies have shown that the increased duration of mitosis in poor carbon is not an artifact caused by poor synchrony (Leitao and Kellogg 2017; Leitao *et al.* 2019). Importantly, we also observed that growth in poor carbon caused a reduction in the maximal extent of hyperphosphorylation reached during mitosis (**Fig. 5 A, B, C**).

**Figure 5:**
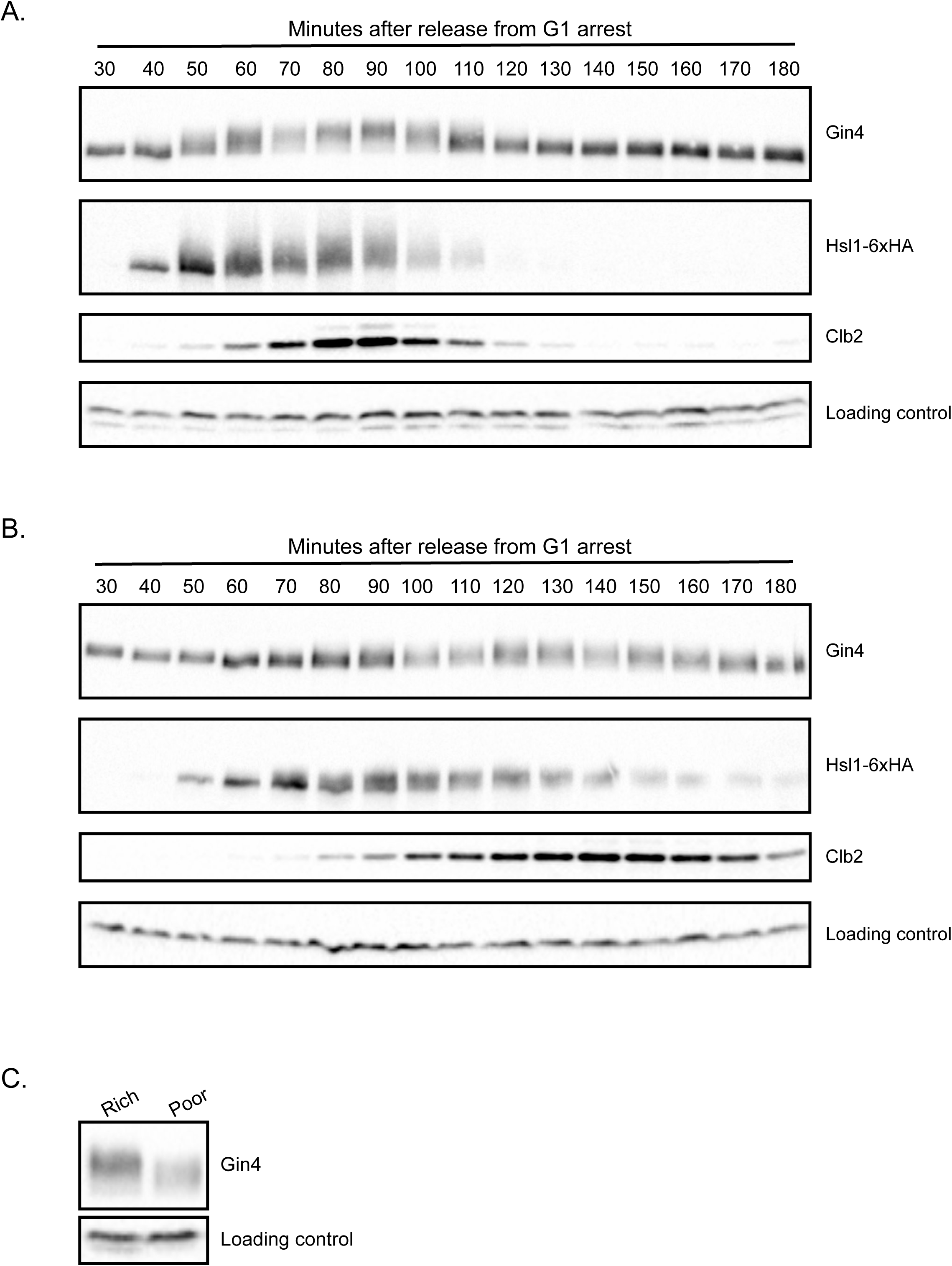
Hyperphosphorylation of Gin4 and Hsl1 is correlated with the extent of bud growth. Wild type cells grown overnight in YPD **(A)** or YPG/E **(B)** were arrested with α factor. The cells were then released from the arrest at 25°C and samples were taken at 10 min intervals. The behavior of Gin4, Hsl1-6XHA and Clb2 was assayed by western blot. **(C)** A direct comparison of the maximal extent of Gin4 phosphorylation in rich or poor carbon was made by comparing samples taken at peak Clb2 expression in each condition (90 min in rich carbon and 140 min in poor carbon). An anti-Nap1 antibody was used as a loading control.

These observations are consistent with the hypothesis that hyperphosphorylation of Gin4 and Hsl1 is correlated with the extent of bud growth rather than a mitotic event that is unrelated to growth. One interpretation of the results is that the extent of hyperphosphorylation of Gin4 and Hsl1 required for mitotic exit is reduced in poor carbon, which allows cells to complete mitosis at a smaller bud size.

### Hyperphosphorylation of Gin4 is dependent upon bud growth

We next tested whether hyperphosphorylation of Gin4 is dependent upon bud growth. To do this, we used a temperature-sensitive allele of *SEC6* (*sec6-4*) to block bud growth. Sec6 is a component of the exocyst complex, which is required at the plasma membrane for docking and fusion of vesicles that drive bud growth. In previous work, we showed that inactivation of Sec6 blocks bud growth and triggers an arrest in early mitosis (Anastasia *et al.* 2012). The arrest is enforced by Swe1. Thus, *sec6-4 swe1*Δ cells fail to undergo bud growth yet enter mitosis and complete chromosome segregation before eventually arresting in late mitosis. We therefore analyzed Gin4 phosphorylation in *sec6-4 swe1*Δ cells, which allowed us to distinguish whether effects of *sec6-4* could be a consequence of a failure to undergo bud growth, or a failure in mitotic progression. As controls, we also analyzed Gin4 hyperphosphorylation in wild type and *swe1*Δ cells.

Cells were released from a G1 arrest and shifted to the restrictive temperature for the *sec6-4* allele before bud emergence. Gin4 phosphorylation was assayed by western blot (**Fig. 6 A**). The same samples were probed for Clb2 as a marker for mitotic progression (**Fig. 6 B**). The *sec6-4 swe1*Δ cells entered mitosis but arrested in late mitosis with high levels of mitotic cyclin, as previously reported (Anastasia *et al.* 2012). Hyperphosphorylation of Gin4 failed to occur in both *sec6-4* and *sec6-4 swe1*Δ cells (**Fig. 6 A, C and Fig. S5 A**). Direct comparison of the extent of Gin4 hyperphosphorylation in mitosis showed a complete loss of Gin4 phosphorylation (**Fig. 6 C**). Thus, hyperphosphorylation of Gin4 is dependent upon membrane trafficking events that drive bud growth. Moreover, since mitotic spindle assembly and chromosome segregation occur normally in *sec6-4 swe1*Δ cells (Anastasia *et al.* 2012), the data further suggest that Gin4 phosphorylation is independent of the signals that drive chromosome segregation.

**Figure 6:**
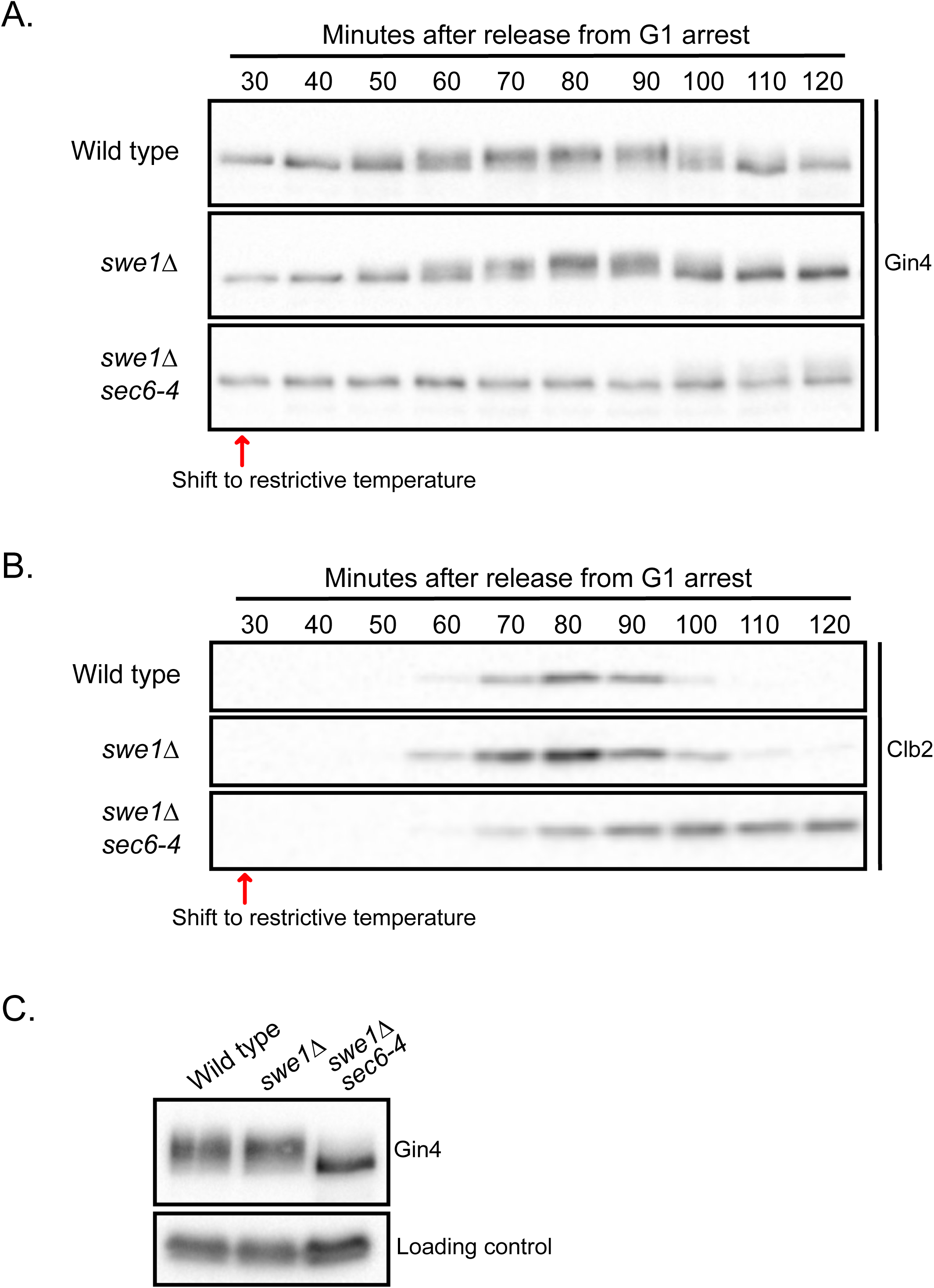
Hyperphosphorylation of Gin4 is dependent upon bud growth. Wild type, *swe1*Δ and *swe1*Δ *sec6-4* cells were released from a G1 arrest in YPD at room temperature and shifted to the restrictive temperature (34°C) 30 min after release from arrest. Samples were taken at the indicated intervals and the behavior of Gin4 **(A)** and Clb2 **(B)** was analyzed by western blot. **(C)** A direct comparison of the extent of Gin4 phosphorylation was made by loading samples from all three strains taken at 90 min (wild type), 80 min (*swe1*Δ) and 100 min (*sec6-4 swe1*Δ).

There is some evidence that growth rate increases with cell size (Jorgensen and Tyers 2004b; Tzur *et al.* 2009; Sung *et al.* 2013). We therefore considered the possibility that Gin4 phosphorylation provides a readout of growth rate rather than a readout of how much growth has occurred. In this case, Gin4 phosphorylation could be driven by dynamic signals generated in association with ongoing events of membrane growth. To investigate, we tested the effects of inactivating Sec6 during bud growth. If Gin4 phosphorylation is a readout of growth rate, an acute block to bud growth should cause rapid loss of Gin4 phosphorylation. In contrast, if Gin4 hyperphosphorylation reports on the extent of bud growth, acute inactivation of Sec6 should stop progression of Gin4 phosphorylation but should not cause a loss of Gin4 phosphorylation. Wild type and *sec6-4* cells were released from a G1 arrest and shifted to the restrictive temperature in early mitosis, as mitotic cyclin levels were rising and Gin4 phosphorylation was beginning to occur. Gin4 phosphorylation persisted in the *sec6-4* cells but did not disappear or increase further after the shift to the restrictive temperature (**Fig. S5 B**). Inactivation of Sec6 late in mitosis when bud growth was largely complete had no effect on Gin4 phosphorylation (**Fig. S5 C**). Thus, Gin4 phosphorylation appears to be correlated with the extent of bud growth, rather than the rate of bud growth. The data further corroborate the model where Gin4 and Hsl1 report on the extent of bud growth until metaphase and do not respond to growth perturbations in anaphase.

### Phosphorylation of Gin4 during bud growth requires binding to anionic phospholipids

We next investigated the mechanisms that drive hyperphosphorylation of Gin4-related kinases, since these could provide clues to how growth-dependent signals are generated and relayed. Both Gin4 and Hsl1 have well-defined C-terminal kinase associated 1 (KA1) domains that bind phosphatidylserine and other anionic phospholipids (Moravcevic *et al.* 2010). Phosphatidylserine is preferentially localized to the growing bud and appears to be the most important effector for KA1 domains (Ejsing *et al.* 2009; Moravcevic *et al.* 2010; Fairn *et al.* 2011b; Klose *et al.* 2012). The KA1 domain of Hsl1 is required for efficient localization to the bud neck, but the mechanism by which the domain promotes bud neck localization remains unknown. More generally, lipid binding domains have been shown to play diverse roles in biochemical mechanisms that generate or relay signals at the plasma membrane, which indicates that their functions reach beyond simply localizing proteins. For example, binding of phosphatidylserine to KA1 domains in protein kinases can promote an open and active conformation that could lead to autophosphorylation or phosphorylation by another kinase (Wu *et al.* 2015; Emptage *et al.* 2017; 2018). Gradual phosphorylation of Gin4 during bud growth is dependent upon Gin4 kinase activity, consistent with a model in which binding of phosphatidylserine drives autophosphorylation (Altman and Kellogg 1997). Together, these observations led us to hypothesize that phosphatidylserine delivered to the plasma membrane during bud growth could drive hyperphosphorylation of Gin4-related kinases, thereby generating a growth-dependent molecular signal that is correlated with the extent of growth. Alternatively, binding to phosphatidylserine could help localize Gin4-related kinases to a location where they can receive and relay growth-dependent signals.

To investigate further, we focused on Gin4. We found that *gin4-*Δ*KA1-GFP* failed to localize to the bud neck normally and was observed primarily in the cytoplasm (**Fig. 7 A**). Weak localization to the bud neck could be detected in a fraction of cells, indicating that determinants outside the KA1 domain contribute to Gin4 localization to the bud neck, as previously seen for *hsl1-*Δ*KA1* (**arrowheads, Fig. 7 A**) (Finnigan *et al.* 2016). We further found that *gin4-*Δ*KA1* cells showed increased cell size and an elongated bud phenotype similar to *gin4*Δ cells (**Fig. 7 B and C**). *gin4-*Δ*KA1* in an *hsl1*Δ background caused a phenotype similar to *gin4*Δ *hsl1*Δ (**Fig. 7 C**). These findings extend results from a previous analysis that utilized an overexpressed version of *gin4-*Δ*KA1-GFP* (Moravcevic *et al.* 2010).

**Figure 7:**
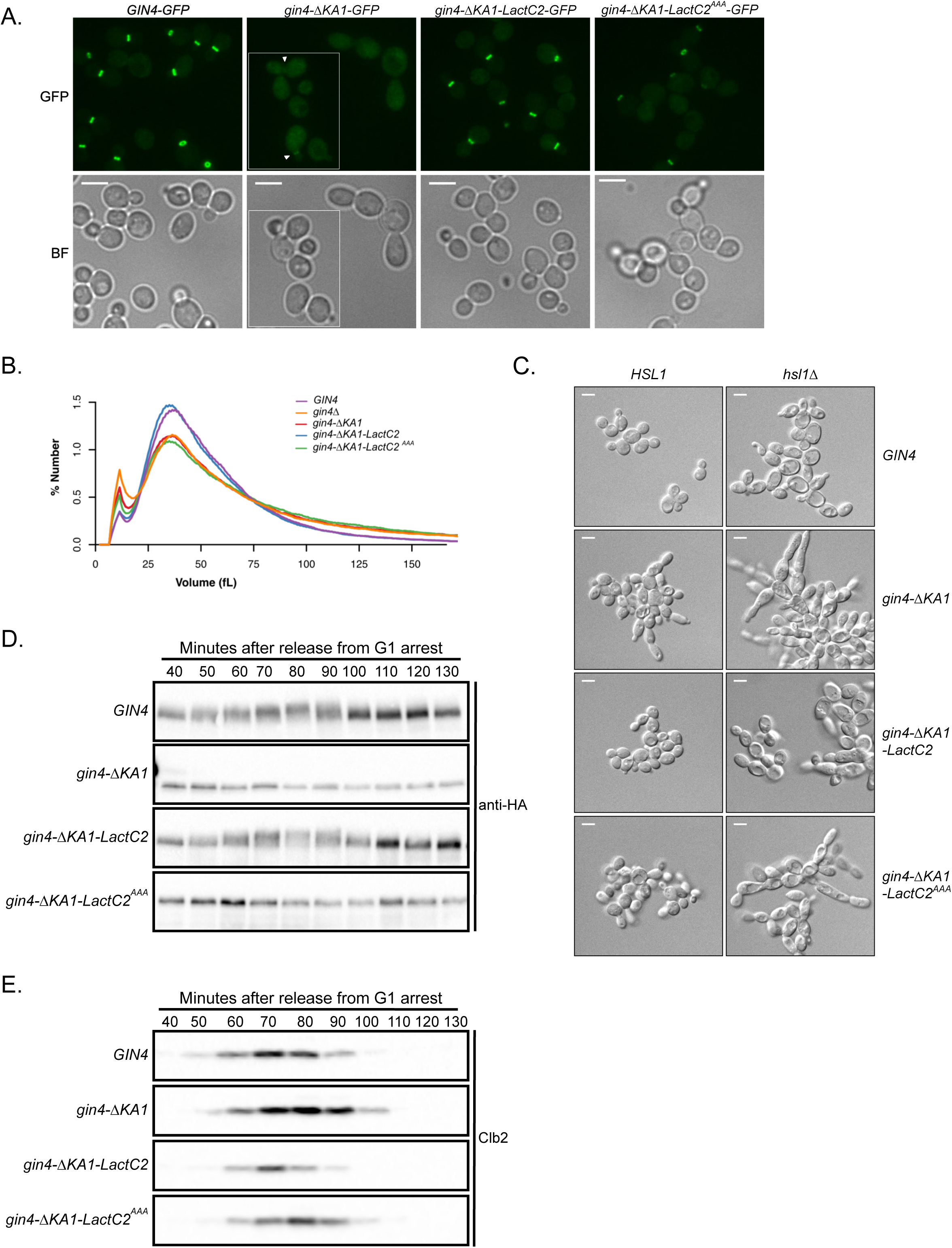
Phosphorylation of Gin4 during bud growth requires binding to anionic phospholipids. **(A**) Cellular localization of Gin4, Gin4-ΔKA1, Gin4-ΔKA1-LactC2 and Gin4-ΔKA1-LactC2^AAA^ fused to GFP at the C-terminus. All four strains were excited by the GFP-laser with identical settings at 100x magnification and displayed with the same brightness levels to compare relative levels of Gin4 localized to the bud neck. Gin4-ΔKA1-GFP shows mostly cytoplasmic localization with a small level of bud neck localization (white arrowheads). The brightfield images for each field are shown below. **(B)** Cells of the indicated genotypes were grown in YPD overnight, diluted in fresh YPD, and then incubated for 5h at 30°C. The size distribution for each strain was analyzed using a Coulter counter. **(C)** Cells of the indicated genotypes were grown to log phase in YPD at 25°C and imaged by DIC optics. **(D and E)** Cells of the indicated genotypes were released from a G1 arrest in YPD at 30°C. The behavior of Gin4 **(D)** and Clb2 **(E)** was analyzed by western blot. In each strain, Gin4 constructs were marked with a 3xHA tag and detected with anti-HA antibody. The signal for *gin4-*Δ*KA1* and the LactC2 constructs was weaker, so these blots were exposed longer. Gin4-ΔKA1-3xHA was about 16 KDa smaller than the other proteins. Scale bar represents 5 µm.

Previous work found that the KA1 domain of Hsl1 could be functionally replaced by a heterologous bovine LactC2 domain that binds phosphatidylserine. Similarly, we found that replacing the KA1 domain of Gin4 with the LactC2 domain rescued the localization defects and mutant phenotype caused by *gin4-*Δ*KA1* (**Fig. 7 A, B, C**). We also found that mutation of three amino acids in the LactC2 domain that are required for efficient binding to phosphatidylserine (*gin4-*Δ*KA1-LactC2^AAA^*) (Yeung *et al.* 2008) caused a phenotype similar to *gin4*Δ and *gin4-*Δ*KA1* (**Fig. 7 A, B, C**). *The* mutant *LactC2^AAA^* domain partially restored Gin4 localization to the bud neck; however, the amount of *gin4-*Δ*KA1-LactC2^AAA^-GFP* at the bud neck was reduced relative to both *GIN4-GFP* and *gin4-*Δ*KA1-LactC2-GFP* (**Fig. 7 A and Fig. S6**). Together, these observations show that the KA1 domain is essential for the function of Gin4, as seen previously for Hsl1 and Kcc4 (Crutchley *et al.* 2009; Moravcevic *et al.* 2010; Finnigan *et al.* 2016).

No previous studies have analyzed whether the KA1 domain influences mitotic duration or the phosphorylation of Gin4-related kinases. We found that *gin4-*Δ*KA1* completely failed to undergo hyperphosphorylation during bud growth (**Fig. 7 D**). Analysis of Clb2 levels in synchronized cells showed that *gin4-*Δ*KA1* cells exhibited an increased duration of mitosis (**Fig. 7 E**). The LactC2 domain was sufficient to restore gradual hyperphosphorylation of Gin4 during bud growth in *gin4-*Δ*KA1-LactC2* cells (**Fig. 7 D**), as well as normal mitotic duration (**Fig. 7 E)**. The mutant version of the LactC2 domain that cannot bind efficiently to phosphatidylserine failed to restore Gin4 hyperphosphorylation (**Figure 7 D**).

Together, these data show that binding to anionic phospholipids is required for gradual hyperphosphorylation and activation of Gin4 during bud growth.

## Discussion

### Gin4-related kinases influence the duration and extent of growth during metaphase

Previous analysis of the functions of Gin4-related kinases used gene deletions. However, phenotypes caused by gene deletions can be the outcome of cumulative defects gained over multiple generations. Therefore, it has not been possible to discern the immediate and direct consequences of loss of function of Gin4-related kinases. Here, conditional alleles allowed us to rigorously define the functions of Gin4-related kinases. We show for the first time that a prolonged delay in metaphase is an immediate consequence of inactivating Gin4 and Hsl1. Growth continues during the delay, leading to aberrant cell size. We also discovered that inactivation of Gin4-related kinases leads to aberrant growth of the mother cell, which indicates that the Gin4-related kinases are required for mechanisms that restrict growth to the daughter bud. Since the Gin4-related kinases are localized to the bud neck, they are ideally positioned to define a domain of growth in the daughter bud. Defects in cytokinesis are another primary consequence of inactivating Gin4-related kinases. Severe defects in mitotic spindle positioning were only observed in the second cell cycle after inactivation of Gin4-related kinases, which suggests that they are an indirect consequence.

The effects of inactivating Gin4 and Hsl1 on the duration and extent of growth in metaphase were not additive. This was surprising because *gin4*Δ and *hsl1*Δ show strong additive effects on cell size and shape (Barral *et al.* 1999). The discovery that loss of Gin4-related kinases causes inappropriate mother cell growth during mitosis suggests an explanation. Previous studies have shown that large mother cells drive increased growth of daughter cells (Schmoller *et al.* 2015; Leitao and Kellogg 2017). Thus, we hypothesize that the increased size of mother cells caused by loss of Gin4 and Hsl1 amplifies aberrant growth in subsequent divisions. However, other factors likely contribute to the additive effects of *gin4*Δ and *hsl1*Δ. For example, the severe spindle positioning defects that appear in the second cell division after inactivation of Gin4 and Hsl1 could cause prolonged mitotic delays that lead to further aberrant growth. Defects in growth control could also be amplified by failures in cytokinesis that create chains of conjoined cells in which the signals that control cell growth and size are no longer effectively compartmentalized.

Our analysis also showed that Gin4 and Hsl1 play different roles in controlling polar bud growth. Destruction of Gin4 caused excessive polar growth in the first cell cycle following destruction, whereas destruction of Hsl1 did not. Previous work suggested that Gin4 binds and regulates Bnr1, a formin protein that controls the location of actin cables that deliver vesicles to sites of membrane growth (Buttery *et al.* 2012). Bnr1 is localized to the bud neck, and loss of Bnr1 is thought to cause inappropriate polar growth because the actin cables that direct isotropic growth are lost (Pruyne *et al.* 2004; Gao *et al.* 2010). Thus, polar growth caused by loss of Gin4 could be due at least partly to misregulation of Bnr1.

Inactivation of Gin4 and Hsl1 had little effect on the duration of anaphase or the extent of growth in anaphase. In a previous study, we found that extensive growth occurs during anaphase, and that the duration of anaphase and the extent of growth in anaphase are both modulated by nutrient-dependent signals (Leitao and Kellogg 2017). Our results suggest that Gin4-related kinases are unlikely to control the anaphase growth interval. The anaphase growth interval is also unlikely to be controlled by Cdk1-inhibitory phosphorylation (Leitao *et al.* 2019). The mechanisms that control anaphase duration in response to nutrient-dependent signals therefore remain mysterious.

### Gin4-related kinases influence the duration of metaphase via Cdk1 inhibitory phosphorylation

The use of conditional alleles allowed us to show that Gin4-related kinases influence metaphase duration, and that they do so primarily via Swe1-dependent Cdk1 inhibitory phosphorylation. Early work suggested that Wee1-related kinases work at mitotic entry. However, more recent work in both vertebrates and yeast found that Wee1 also controls events after mitotic entry (Deibler and Kirschner 2010; Harvey *et al.* 2011; Lianga *et al.* 2013; Vassilopoulos *et al.* 2014; Toledo *et al.* 2015; Leitao *et al.* 2019). Several previous studies implied that Gin4-related kinases could be responsible for controlling Cdk1 inhibitory phosphorylation during metaphase; however, interpretation of the results was complicated by the use of gene deletions, and the experiments did not directly test whether Gin4-related kinases control metaphase via Cdk1 inhibitory phosphorylation (Carroll *et al.* 1998; Barral *et al.* 1999; Sreenivasan and Kellogg 1999; Sreenivasan *et al.* 2003). Here, acute conditional inactivation of Gin4 and Hsl1 provided definitive evidence that Gin4-related kinases influence the duration of metaphase via Cdk1 inhibitory phosphorylation. We also showed that loss of Gin4-related kinases causes a failure in full hyperphosphorylation of Swe1, which is thought to be required for inactivation of Swe1. Previous studies have shown that Gin4-related kinases from fission yeast can directly phosphorylate Wee1; however, there is no evidence yet that this is true in budding yeast (Coleman *et al.* 1993; Kanoh and Russell 1998; Opalko *et al.* 2019).

Analysis of the effects of conditional inactivation of Gin4-related kinases suggested that they do not influence the duration of metaphase solely via Swe1. Previous studies showed that the duration of metaphase in *swe1*Δ cells is shorter than wild type cells (Lianga *et al.* 2013; Leitao *et al.* 2019). Here, we found that *swe1*Δ substantially reduces the metaphase delay caused by inactivation of Gin4-related kinases but did not cause metaphase duration to be shorter than wild type cells. Moreover, *swe1*Δ did not fully rescue the elongated buds, cell separation defects and inappropriate mother cell growth caused by loss of Gin4-related kinases. Together, these observations suggest that Gin4-related kinases influence growth in metaphase partly via a mechanism that works downstream or independently of Cdk1 inhibitory phosphorylation. The fission yeast homolog of Gin4 is also known to execute functions that are independent of Wee1 (Breeding *et al.* 1998).

### Hyperphosphorylation of Gin4-related kinases is dependent upon bud growth

Gin4 and Hsl1 undergo gradual hyperphosphorylation and activation during mitosis. Growth of the bud also occurs gradually during mitosis, and loss of Gin4-related kinases causes inappropriate growth during a metaphase delay. These and other data led us to hypothesize that hyperphosphorylation of Gin4-related kinases reflects the activity of signaling mechanisms that measure bud growth during mitosis. To begin to test this hypothesis, we further investigated the correlation between bud growth and hyperphosphorylation of Gin4 and Hsl1. To do this, we analyzed hyperphosphorylation of Gin4 and Hsl1 in cells growing in poor carbon, which increases the duration of bud growth in mitosis, while also decreasing the extent of growth. Poor carbon prolonged the interval during which hyperphosphorylation of Gin4 and Hsl1 took place in mitosis. Importantly, poor carbon also reduced the maximal extent of Gin4 and Hsl1 hyperphosphorylation achieved in mitosis. Thus, hyperphosphorylation of Gin4 and Hsl1 is correlated with both the duration and extent of growth in mitosis.

The increased duration of Gin4 and Hsl1 hyperphosphorylation in poor carbon is unlikely to be due to poor cell cycle synchrony because analysis of mitotic spindle dynamics in both live single cells and synchronized populations of cells has shown that the durations of both metaphase and anaphase are substantially increased in poor carbon (Leitao and Kellogg 2017; Leitao *et al.* 2019). Moreover, poor synchrony cannot explain why the maximal extent of Gin4 and Hsl1 hyperphosphorylation is dramatically reduced in poor carbon. If poor synchrony were the cause, one would expect to see a lower fraction of Gin4 reaching the maximally hyperphosphorylated state in poor carbon. Rather, we observe a complete loss of the maximally hyperphosphorylated forms of Gin4 and Hsl1 in poor carbon.

To further investigate the relationship between bud growth and phosphorylation of Gin4 we tested whether hyperphosphorylation of Gin4 is dependent upon growth. We found that blocking membrane trafficking events that drive plasma membrane growth causes a complete failure in Gin4 phosphorylation. Furthermore, blocking membrane growth after bud growth was initiated halted the progression of Gin4 phosphorylation, but did not cause a loss of Gin4 phosphorylation, consistent with the idea that the Gin4 phosphorylation is correlated with the extent of bud growth, rather than growth rate or time spent in mitosis.

Blocking membrane growth causes a prolonged metaphase delay that is dependent on Cdk1 inhibitory phosphorylation (Anastasia *et al.* 2012). Elimination of Swe1-mediated Cdk1 inhibitory phosphorylation abrogates the delay but does not restore Gin4 hyperphosphorylation, suggesting that Gin4 hyperphosphorylation is upstream of Cdk1 inhibitory phosphorylation. Moreover, in previous work we found that Gin4 hyperphosphorylation occurs normally in cells treated with drugs that block the formation of a mitotic spindle (Mortensen *et al.* 2002). Thus, Gin4 hyperphosphorylation is dependent upon membrane trafficking events that drive bud growth and appears to be independent of the events of mitotic spindle assembly and chromosome segregation.

### Hyperphosphorylation of Gin4 requires binding to anionic phospholipids

We found that growth-dependent hyperphosphorylation of Gin4 is dependent upon the KA1 domain, which binds anionic phospholipids. Moreover, the functions of the KA1 domain that are required for Gin4 phosphorylation could be fully replaced by a heterologous bovine LactC2 domain that binds phosphatidylserine. Mutations in the LactC2 domain that reduce binding to phosphatidylserine block hyperphosphorylation of Gin4 and cause a phenotype similar to *gin4*Δ. Together, these observations demonstrate that gradual hyperphosphorylation of Gin4 during bud growth is dependent upon binding to anionic phospholipids.

The KA1 domain binds preferentially to phosphatidylserine but can also bind other anionic phospholipids, such as phosphatidylinositol (Moravcevic *et al.* 2010; Wu *et al.* 2015). In contrast, the LactC2 domain appears to bind only to phosphatidylserine (Shao *et al.* 2008). The fact that the KA1 domain can be functionally replaced by the LactC2 domain therefore suggests that binding to phosphatidylserine is sufficient to drive gradual hyperphosphorylation of Gin4 (Moravcevic *et al.* 2010; Fairn *et al.* 2011a). Phosphatidylserine is preferentially localized to the growing bud and is enriched at the bud neck (Moravcevic *et al.* 2010; Fairn *et al.* 2011a). Phosphatidylserine is also preferentially localized to sites of membrane growth in fission yeast (Haupt and Minc 2017).

Hyperphosphorylation of both Gin4 and Hsl1 is dependent upon their kinase activity, which suggests that gradual hyperphosphorylation of these kinases during bud growth is due to autophosphorylation (Altman and Kellogg 1997; Barral *et al.* 1999). Moreover, previous studies suggested that binding of anionic phospholipids to KA1 domains in protein kinases that are related to Gin4 can drive formation of an open, active conformation (Wu *et al.* 2015; Emptage *et al.* 2017; 2018). Together, these observations suggest that anionic phospholipids delivered to the growing bud could drive autophosphorylation by directly binding and activating Gin4-related kinases. This model could explain why *gin4-*Δ*KA1-LactC2^AAA^* causes a complete loss of Gin4 phosphorylation, despite allowing a fraction of Gin4 to be localized to the bud neck. Alternatively, binding of Gin4-related kinases to anionic phospholipids could help recruit them to a location where they receive other signals that are required for hyperphosphorylation. Both scenarios could help generate or relay a molecular readout of the extent of bud growth.

### A growth-dependent signaling hypothesis for the functions of Gin4-related kinases

Together, the data suggest a model in which anionic phospholipids delivered to the growing bud bind and activate Gin4-related kinases, thereby generating a signal that is correlated with the extent of growth during metaphase. We further propose that once the signal reaches a threshold, Gin4-related kinases drive full hyperphosphorylation and inactivation of Swe1, thereby triggering mitotic progression and termination of the metaphase growth interval. We refer to this model as the growth-dependent signaling hypothesis.

The data do not yet rule out alternative models. Nevertheless, the growth-dependent signaling hypothesis is consistent with the available data. It would explain why loss of Gin4-related kinases causes an abnormally prolonged interval of growth during metaphase, leading to aberrant cell size. In this case, the hypothesis suggests that loss of Gin4-related kinases causes cells to behave as though they cannot measure how much bud growth has occurred. The hypothesis also explains why Gin4-related kinases undergo gradual hyperphosphorylation and activation during bud growth, and why hyperphosphorylation is dependent upon membrane trafficking events that drive cell growth yet seemingly independent of the events of mitotic spindle assembly and chromosome segregation. In this model, the Gin4-related kinases would be direct molecular sensors of a critical event that drives cell growth (i.e. delivery of anionic phospholipids to the plasma membrane). The data are also consistent with a model in which the Gin4-related kinases relay a growth-dependent signal generated by another mechanism.

The growth-dependent signaling hypothesis could also explain why the maximal extent of Gin4 and Hsl1 hyperphosphorylation achieved in mitosis appears to be reduced in poor nutrients. We hypothesize that the reduction in cell size at completion of metaphase in poor nutrients is driven by nutrient-dependent signals that reduce the threshold activity of Gin4-related kinases required for mitotic progression.

A key question concerns the cause of the prolonged metaphase delay observed in cells that lack Gin4-related kinases. We favor a model in which the delay is a direct consequence of a failure in growth-dependent signals that are used to detect and measure cell growth. However, the data do not yet rule out the possibility that a more proximal cause of the delay is failure of an event that is unrelated to cell growth.

Measuring cell growth cannot be the sole function of Gin4-related kinases. Gin4-related kinases are required for organization of septins at the bud neck, and for controlling the pattern and location of growth. Gin4-related kinases also appear to be embedded in the TORC2 network, which influences cell size and growth rate, although the functional relationships between the TORC2 network and Gin4-related kinases are poorly understood (Alcaide-Gavilan *et al.* 2018). In each case, Gin4-related kinases appear to be closely associated with signals that influence the location, pattern and extent of growth. Thus, it is possible that the mechanisms that measure growth are embedded in the mechanisms that drive and coordinate the events of cell growth.

Theoretical analysis has shown that cell size control can be achieved by an “adder” mechanism, in which a constant increment of growth is added during each cell cycle (Campos *et al.* 2014). In the adder model, cells measure growth, rather than size. Adder behavior has been reported in cells ranging from bacteria to vertebrates, yet a mechanistic explanation for how growth could be measured has remained elusive (Campos *et al.* 2014; Cadart *et al.* 2018). Here, we propose that growth-dependent activation of Gin4-related kinases events could be part of an adder mechanism that measures bud growth. Thus, delivery of signaling lipids to sites of growth could be the critical event that is monitored to measure bud growth. Growth-dependent signaling could be broadly relevant, as it would be readily adaptable to cells of diverse size and shape. It could also influence cell shape by controlling the extent of growth at specific locations on the cell surface. Further analysis of the mechanisms that drive growth-dependent signaling should yield new insights into control of cell growth and size.

## Materials and Methods

### Yeast strain construction, media, and reagents

All strains are in the W303 background (*leu2-3,112 ura3-1 can1-100 ade2-1 his3-11,15 trp1-1 GAL+ ssd1-d2*). The additional genetic features of strains are listed in Table S1. Cells were grown in YP medium (1% yeast extract, 2% peptone, 40 mg/liter adenine) supplemented with 2% dextrose (YPD), or 2% glycerol and 2% ethanol (YPG/E). For live cell imaging, cells were grown in complete synthetic medium (CSM) supplemented with 2% dextrose and 40 mg/ml adenine.

Gene deletions and C-terminal epitope tagging was performed by standard PCR amplification and homologous recombination (Longtine *et al.* 1998; Janke *et al.* 2004; Lee *et al.* 2013). *gin4-*Δ*KA1-LactC2* constructs integrated at the GIN4 locus were created by gene splicing with overlap extension (Horton *et al.* 1990). Briefly, LactC2 fragments were PCR amplified from the plasmids pKT2100 or pKT1995 (Takeda *et al.* 2014) with 40bp of flanking sequence at the 5’ end that was homologous to the GIN4 ORF just upstream of the KA1 domain (amino acids 1007-1142) using oligos Gin4-39 and Gin4-40 (Table 2). The 3xHA::His3MX6 fragment was amplified from pFA6a-3HA-His3MX6 (Longtine *et al.* 1998) with 5’ homology to the terminal sequence of LactC2 and 3’ homology to the DNA sequence just downstream of the Gin4 ORF using primers Gin4-38 and Gin4-41. The two fragments were then gel purified, annealed to each other and elongated for 15 PCR cycles in the absence of primers, followed by PCR amplification using primers Gin4-38 and Gin4-39. The resulting fragments were then transformed into wild type cells and correct integrants were identified by western blotting with anti-HA antibody. To create the GFP-tagged versions of the *gin4-*Δ*KA1-LactC2* constructs, GFP-His3MX6 was amplified from pFA6a-GFP-His3MX6 (Longtine *et al.* 1998) and spliced to LactC2 as described above.

**Table 1:** Strains used in this study.

| <u>Strain</u> | <u>Mating type</u> | <u>Genotype</u> | <u>Source</u> |
| --- | --- | --- | --- |
| DK 186 | a | *bar1*Δ | (Altman and Kellogg 1997) |
| DK 3418 | a | *bar1*Δ *HSL1-6XHA::His3MX6* | This study |
| SH 24 | a | *bar1*Δ *swe1*Δ::URA3 | (Harvey and Kellogg 2003) |
| DK 1600 | a | *bar1*Δ *swe1*Δ::His3MX6 *sec6-4::KanMX6* | (Anastasia et al. 2012) |
| DK 3510 | a | *bar1*Δ *his3::His3MX6+TIR1 leu2::LEU2+TIR1 SPC42-yomRuby2::KanMX6* | This study |
| DK 3307 | a | *bar1*Δ *his3::His3MX6+TIR1 leu2::LEU2+TIR1 SPC42-GFP::HphNTI gin4-AID::KanMX6* | This study |
| DK 3308 | a | *bar1*Δ *his3::His3MX6+TIR1 leu2::LEU2+TIR1 SPC42-GFP::HphNTI hsl1-AID::TRP* | This study |
| DK 3327 | a | *bar1*Δ *his3::His3MX6+TIR1 leu2::LEU2+TIR1 SPC42-GFP::HphNTI gin4-AID::KanMX6 hsl1-AID::TRP1* | This study |
| DK 3330 | a | *bar1*Δ *his3::His3MX6+TIR1 leu2::LEU2+TIR1 SPC42-GFP::HphNTI gin4-AID::KanMX6 hsl1-AID::TRP1 swe1*Δ::URA3 | This study |
| DK 3350 | a | *bar1*Δ *GIN4-GFP:: His3MX6 SPC42-yomRuby2::KanMX* | This study |
| DK 3790 | a | *bar1*Δ *gin4-ΔKA1-GFP::His3MX6 SPC42-yomRuby2::KanMX* | This study |
| DK 3351 | a | *bar1*Δ *gin4-ΔKA1-LactC2-GFP:: His3MX6 SPC42-yomRuby2::KanMX* | This study |
| DK 3823 | a | *bar1*Δ *gin4-ΔKA1-LactC2<sup>AAA</sup>-GFP::His3MX6 SPC42-yomRuby2::KanMX* | This study |
| DK 888 | a | *bar1*Δ *gin4*Δ::LEU2 | (Mortensen et al. 2002) |
| HT159 | a | *bar1*Δ *hsl1*Δ::His3MX6 | This study |
| DK3784 | a | *bar1Δ gin4Δ::LEU2 hsl1Δ::His3MX6* | This study |
| DK2158 | a | *bar1Δ gin4Δ::LEU2 hsl1Δ::HphNTI swe1Δ::URA3* | This study |
| DK 373 | a | *bar1Δ GIN4-3xHA::TRP1* | This study |
| DK 2822 | a | *bar1Δ gin4-ΔKA1-3xHA::His3MX6* | This study |
| DK 3286 | a | *bar1Δ gin4-ΔKA1-LactC2-3xHA:: His3MX6* | This study |
| DK 3295 | a | *bar1Δ gin4-ΔKA1-LactC2<sup>AAA</sup>-3xHA::His3MX6* | This study |
| DK3621 | a | *bar1Δ GIN4-3xHA::TRP1 hsl1Δ::HphNTI* | This study |
| DK3624 | a | *bar1Δ gin4-ΔKA1-3xHA::His3MX6 hsl1Δ::HphNTI* | This study |
| DK3627 | a | *bar1Δ gin4-ΔKA1-LactC2-3xHA:: His3MX6*<br>*hsl1Δ::HphNTI* | This study |
| DK3630 | a | *bar1Δ gin4-ΔKA1-LactC2<sup>AAA</sup>-3xHA::His3MX6*<br>*hsl1Δ::HphNTI* | This study |
| DK1440 | a | *bar1Δ sec6-4::KanMX6* | (Anastasia et al. 2012) |

**Table 2:**
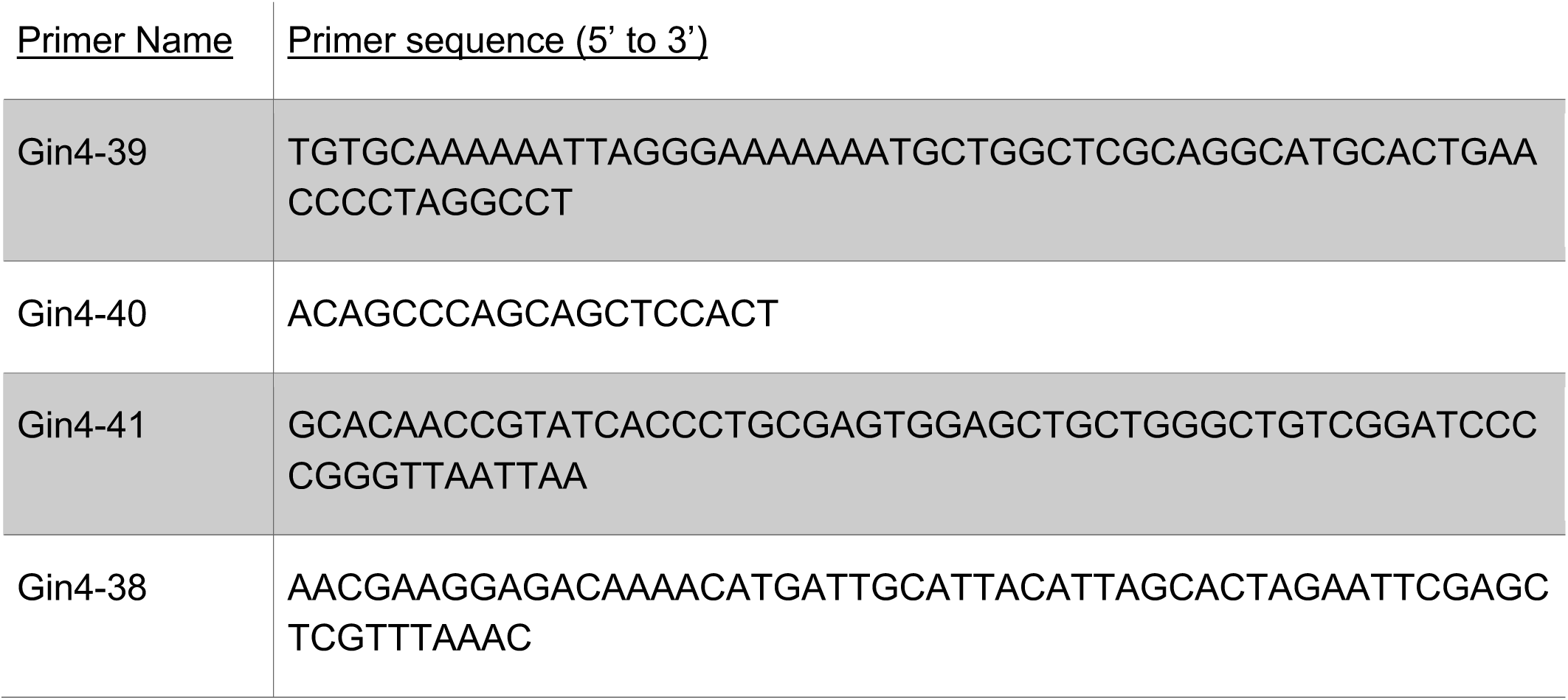
Primers used in this study.

To generate strains with an AID tag on *GIN4* and/or *HSL1*, the *HSL1* gene was tagged at the C-terminus with an AID tag marked with KanMX6 in a parent strain that has two copies of the *TIR1* gene. The KanMX6 marker was then replaced by a *TRP1* marker. Next, a second AID tag marked with KanMX6 was incorporated at the *GIN4* locus. The *SPC42* gene in all four AID-tagged strains was fused to GFP at the C-terminus using standard PCR and homologous recombination. The parent strain that contains *2xTIR1* was used as the control strain and was modified to express endogenous SPC42 fused with yeast-optimized mRuby2 (yomRuby2). Auxin was dissolved in 100% ethanol to make a 50 mM stock solution.

### Cell cycle time courses and Western blotting

Cell cycle time courses were carried out as previously described (Harvey *et al.* 2011). Briefly, cells were grown to log phase at room temperature overnight in YPD or YPG/E to an optical density (OD_600_) of 0.5 - 0.7. Cultures were adjusted to the same optical density and were then arrested in G1 phase by incubation in the presence of 0.5 µg/mL α factor at room temperature for 3 hours. Cells were released from the arrest by washing 3 times with fresh YPD or YPG/E. All time courses were carried out at 25°C unless otherwise noted, and α factor was added back at 70 minutes to prevent initiation of a second cell cycle. For experiments involving auxin-mediated destruction of proteins, a single culture synchronized in G1 phase was split into two culture flasks and 0.5 mM auxin was added to one flask at 20 minutes after release from the G1 phase arrest. An equivalent volume of ethanol was added to the control flask.

For western blotting, 1.6 mL sample volumes were collected in screw cap tubes and centrifuged at 13,000 rpm for 30 sec. After discarding the supernatant, 200 µL acid washed glass beads were added to the tubes and the samples were frozen in liquid nitrogen. Cells were lysed in 140 µL sample buffer (65 mM Tris HCl, pH-6.8, 3% SDS, 10% glycerol, 50 mM sodium fluoride, 100 mM ß-glycerophosphate, 5% ß-mercaptoethanol, and bromophenol blue) supplemented with 2 mM PMSF immediately before use. For experiments involving immunoblotting for Gin4-AID proteins, the sample buffer also included the protease cocktail LPC (1 mg/mL leupeptin, 1 mg/mL pepstatin, 1 mg/mL chymostatin dissolved in dimethylsulfoxide; used at 1/500 dilution). Sample buffer was added to cells immediately after they were removed from liquid nitrogen and the cells were then lysed in a Mini-beadbeater 16 (BioSpec) at top speed for 2 min. After a brief centrifugation, the samples were placed in a boiling water bath for 5 min and were then centrifuged again at 13,000 rpm for 3 min before loading onto SDS-PAGE gels. SDS PAGE was carried out as previously described (Harvey *et al.* 2011). 10% polyacrylamide gels with 0.13% bis-acrylamide were used for analysis of Gin4, Clb2, and Nap1 (loading control). 9% polyacrylamide gels with 0.14% bis-acrylamide were used for Hsl1 and Swe1 blots. Proteins were immobilized onto nitrocellulose membranes using wet transfers for 1h 45 min. Blots were probed with the primary antibody at 1-2 µg/mL at room temperature overnight in 5% milk in PBST (1x phosphate buffered saline, 250 mM NaCl, 0.1% Tween-20) with 0.02% sodium azide. All the primary antibodies used in this study are rabbit polyclonal antibodies generated as described previously (Kellogg and Murray 1995; Sreenivasan and Kellogg 1999; Mortensen *et al.* 2002). Primary antibodies were detected by an HRP-conjugated donkey anti-rabbit secondary antibody (GE Healthcare; # NA934V) incubated in PBST for 1h at room temperature. Blots were rinsed in PBS before detection via chemiluminescence using ECL reagents (Advansta; #K-12045-D50) with a Bio-Rad ChemiDoc imaging system.

### Coulter counter analysis

Cell cultures were grown in 10 mL YPD medium to an OD_600_ between 0.4 - 0.6. Cells were fixed by addition of 1/10 volume of 37% formaldehyde to the culture medium followed by incubation at room temperature for 1h. Cells were then pelleted and resuspended in 0.5 mL PBS containing 0.02% sodium azide and 0.1% Tween-20 and analyzed on the same day. Cell size was measured using a Coulter counter (Channelizer Z2; Beckman Coulter) as previously described (Jorgensen *et al.* 2002; Artiles *et al.* 2009). Briefly, 40 μL of fixed cells were diluted in 10 mL diluent (Isoton II; Beckman Coulter) and sonicated for 5 pulses of approximately 0.5 second each at low power. The Coulter Counter data shown in the figures represents the average of 3 biological replicates that is each the average of 3 technical replicates. For **Fig. 7 B** the strains were grown to log phase overnight at room temperature, diluted to OD_600_ - 0.1 in 5 mL fresh YPD, and then incubated for 4-5 h at 30°C to observe temperature-dependent phenotypes of the mutants.

### Microscopy

For DIC imaging, cells were grown to log phase in YPD and fixed in 3.7% formaldehyde for 30 min and then resuspended in PBS with 0.1% Tween-20 and 0.02% sodium azide. Images were obtained using a Zeiss-Axioskop 2 Plus microscope fitted with a 63x Plan-Apochromat 1.4 n.a. objective and an AxioCam HR camera (Carl Zeiss, Thornwood, NY). Images were acquired using AxioVision software and processed on Fiji (Schindelin *et al.* 2012).

For live cell time-lapse imaging, the control strain (DK3510) and AID-tagged strains (DK3307, DK3308, DK3327 or DK3330) were grown in CSM overnight to an OD_600_ of 0.1 - 0.2 and then arrested in G1 phase with α factor. The control and the AID-tagged strains were mixed in a 1.6 mL tube and then washed 3X in CSM prewarmed to 30°C to release the cells from the G1 phase arrest. After resuspending the cells in CSM, approximately 200 µL cells were immobilized onto a concanavalin A-treated chambered #1.5 Coverglass system (Labtek-II; Nunc^(tm)^#155409) for 5 min. Unbound cells were washed away by repeated washes with CSM. The cells were then incubated in 500 µL CSM at 27°C for the duration of the imaging. Auxin was added to the cells to a final concentration of 0.5 mM 20 minutes after the first wash used to release the cells from the α factor arrest.

Scanning confocal images were acquired on a Zeiss 880 confocal microscope running ZEN Black software using a 63x/1.4 n.a. Plan Apo objective. The microscope was equipped with a heat block stage insert with a closed lid and exterior chamber for temperature control. The microscope was allowed to equilibrate at the set temperature of 27°C for at least 1h to ensure temperature stability prior to imaging. Definite Focus was used to keep the sample in focus during the duration of the experiment. 1 x 2 tiled *z*-stack images were acquired every 3 min. Zoom and frame size were set to 0.8x magnification to achieve a consistent pixel area of 1024 x 1024 pixels in XY and pixel dwell time was 0.5 µs. Optical sections were taken for a total of 14 *z*-planes every μm with frame averaging set to 2, to reduce noise. 488 nm laser power was set to 0.2 % and the 561 nm laser power was set to 1% to minimize cell damage. The gain for GFP, RFP and bright field was set to 550, 750 and 325, respectively. The same gain settings were used for each experiment. GFP signal was acquired on a GaAsP detector and collected between 498 nm - 548 nm. Brightfield images were collected simultaneously. RFP signal was acquired on a GaAsP detector and collected between 577 nm - 629 nm.

Initiation of metaphase is defined as the time at which the spindle poles move to a distance of 1-2 µM apart in the mother cell (Leitao and Kellogg 2017). Initiation of anaphase is marked by further separation of spindle poles and migration of one spindle pole into the daughter cell. Completion of anaphase correspond to the time when the spindle poles reach their maximal distance apart.

To visualize the localization of Gin4 constructs fused to GFP, cells were grown in CSM overnight, fixed in 3.7% formaldehyde for 15 min, and then resuspended in 500 µL 1x PBST. Images were acquired on a spinning disk confocal microscope with a Solamere system running MicroManager* (Schindelin *et al.* 2012). The microscope was based on a Nikon TE2000 stand and Coherent OBIS lasers. We used a 100x/1.4 n.a. Plan Apo objective for **Fig. 7 A** and a 63x/1.4 Plan Apo objective for data collection in **Fig. S6**. Pixel sizes were 0.11 µm in X,Y and *z*-stack spacing was set to 0.5 µm with a total of 17 *z*-slices. GFP was excited at 488 nm and collected through a 525/50nm band pass filter (Chroma) onto a Hamamatsu ImageEMX2 EMCCD camera. Gain levels were set to 200 to maximize signal without hitting saturation. GFP and brightfield images were collected sequentially.

### Image analysis

All images were analyzed on Fiji (Schindelin *et al.* 2012). For visualization of GFP-tagged Gin4 constructs, a sum projection of *z*-slices was used. Movies for the time-lapse were processed as previously described (Ferrezuelo *et al.* 2012). The bright field images were processed using the “Find Focused Slices” plugin available on Fiji to create a stack with the focused slice +/- one slice for each timepoint. A *z*-projection with sum of slices was performed on this stack and then bud volumes were determined using the plugin BudJ (Ferrezuelo *et al.* 2012).

The timings of cell cycle events were determined as previously reported (Altman and Kellogg 1997). Briefly, bud initiation was manually determined by the appearance of a protrusion on the surface of the mother cell. The initiation of metaphase was marked by the appearance of separation of spindle poles to 2-3 microns apart. Initiation of anaphase was marked by further separation of the spindle poles and segregation of one of the poles into the daughter cell. We defined completion of anaphase as the point at which the spindle poles reached their maximal distance apart.

For a quantitative comparison of the localization of GFP-tagged Gin4 constructs in **Fig. S6**, a *z-*projection with sum of slices was performed on the images and an elliptical ROI was drawn around the bud neck. The maximum pixel intensity was determined for each cell after subtracting the background pixel intensity.

### Statistical analysis

Data acquired from the image analysis were plotted as scatter dot plots using GraphPad Prism. The scatter plots show the data distribution along with the mean and standard deviation for each strain. For all scatter dot plots, the unpaired t-test was calculated using the Mann-Whitney test for non-Gaussian distributions and the two-tailed *p-*values have been mentioned.

### Online supplemental material

Fig. S1 relates to Fig. 1 and provides a characterization of the *gin4-AID* and *hsl1-AID alleles.* Fig. S2 relates to Fig. 2 and shows further evidence for the role of Gin4 and Hsl1 in the control of bud growth. Fig. S3 relates to Fig. 3 and shows a time-dependent increase in the *gin4-AID hsl1-AID* phenotype. Fig. S4 relates to Fig. 4 and shows the rapid response of Swe1 dephosphorylation upon the inactivation of Gin4 and Hsl1. Fig. S5 relates to Fig. 6 and shows the effect of growth inactivation at different times during bud growth on Gin4 phosphorylation. Fig. S6 relates to Fig. 7 and shows peak pixel intensity of the various Gin4-GFP constructs localized to the bud neck. Video 1 and Video 2 show time-lapse videos of *2xTIR1* and *gin4-AID hsl1-AID* cells as they progress through the cell cycle. The tables show yeast strains (Table 1), primers (Table 2), and plasmids (Table 3) used in this study.

**Table 3:** Plasmids used in this study.

| **<u>Plasmid Name</u>** | **<u>Details</u>** | **<u>Source</u>** |
| --- | --- | --- |
| pKT1995 | pRS306-GFP-LactC2<sup>AAA</sup>-URA3 | (Takeda *et al.* 2014) |
| pKT2100 | pRS306-GFP-LactC2-URA3 | (Takeda *et al.* 2014) |
| pAID1 | AID-tagging genes at the C-terminus;<br>Kanamycin resistance | (Nishimura *et al.* 2009) |
| pTIR2 | Plasmid containing the osTIR1 under the<br>GPD1 promoter. After PmeI digestion,<br>recombines at the *HIS3* locus | (Nishimura *et al.* 2009) |
| pTIR4 | Plasmid containing the osTIR1 under the<br>GPD1 promoter. After PmeI digestion,<br>recombines at the *LEU2* locus | (Nishimura *et al.* 2009) |
| pFA6a-<br>yomRuby2::KanMx | Yeast-optimized (yo) mRuby2 | (Lee *et al.* 2013) |

## Data availability

Strains and plasmids are available upon request. The authors affirm that all data necessary for confirming the conclusions of the article are present within the article, figures, and tables.

## Acknowledgements

We thank Ben Abrams and the UCSC Life Microscopy Facility for assistance with microscopy. The authors declare no competing financial interests.

## Figure legends

**Figure S1:**
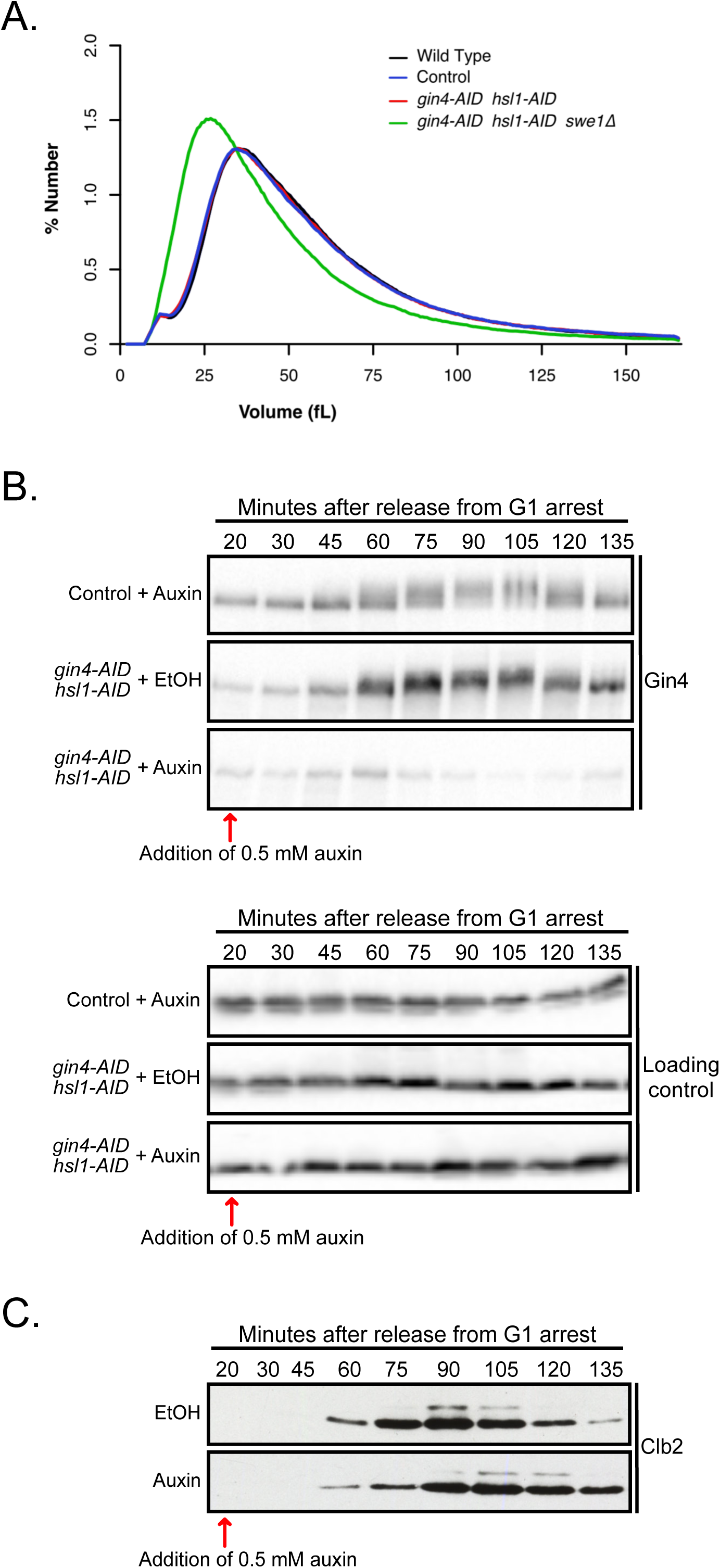
Characterization of *gin4-AID* and *hsl1-AID* alleles. **(A)** Wild type, *2xTIR1* control cells, *gin4-AID hsl1-AID 2xTIR1 cells* and *gin4-AID hsl1-AID swe1*Δ *2xTIR1* cells were grown overnight at room temperature in YPD and cell size distributions were analyzed with a Coulter counter. **(B)** Control cells and *gin4-AID hsl1-AID* cells growing in YPD were released from a G1 phase arrest. After release, the *gin4-AID hsl1-AID* cells were split into two aliquots and 0.5 mM auxin was added to one aliquot and an equivalent amount of the solvent for auxin was added to the other. Auxin was also added to the control strain. Samples were taken at the indicated intervals and the behavior of Gin4 was analyzed by western blot. All strains contain 2 copies of the *TIR1* gene (*2xTIR1*). **(C)** *gin4-AID hsl1-AID 2xTIR1* cells growing in YPD were released from a G1 phase arrest. After release, the cells were split into two aliquots and 0.5 mM auxin was added to one aliquot and an equivalent amount of the solvent for auxin was added to the other. Samples were taken at the indicated intervals and the behavior of Clb2 was analyzed by western blot.

**Figure S2:**
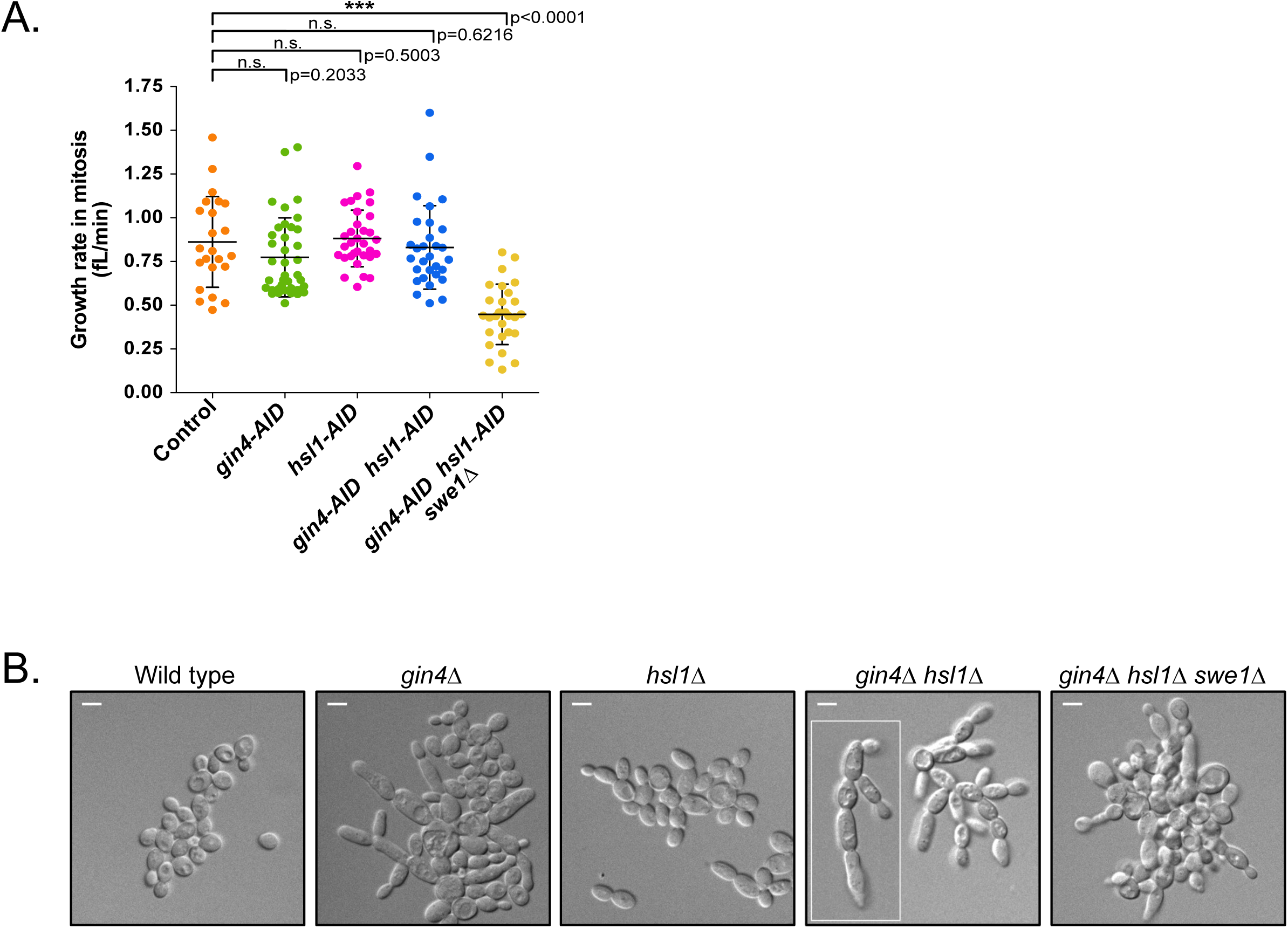
Gin4 and Hsl1 control bud growth during mitosis. **(A)** A scatter plot showing the growth rate of the daughter buds during mitosis for the indicated genotypes. The growth rate (fL/min) was determined as the increase in bud volume from initiation of metaphase to the completion of anaphase, divided by the total time spent in metaphase and anaphase. **(B**) *gin4*Δ, *hsl1*Δ, *gin4*Δ *hsl1*Δ, *and gin4*Δ *hsl1*Δ *swe1*Δ cells were grown to log phase in YPD media at 25°C and images were obtained using DIC optics.

**Figure S3:**
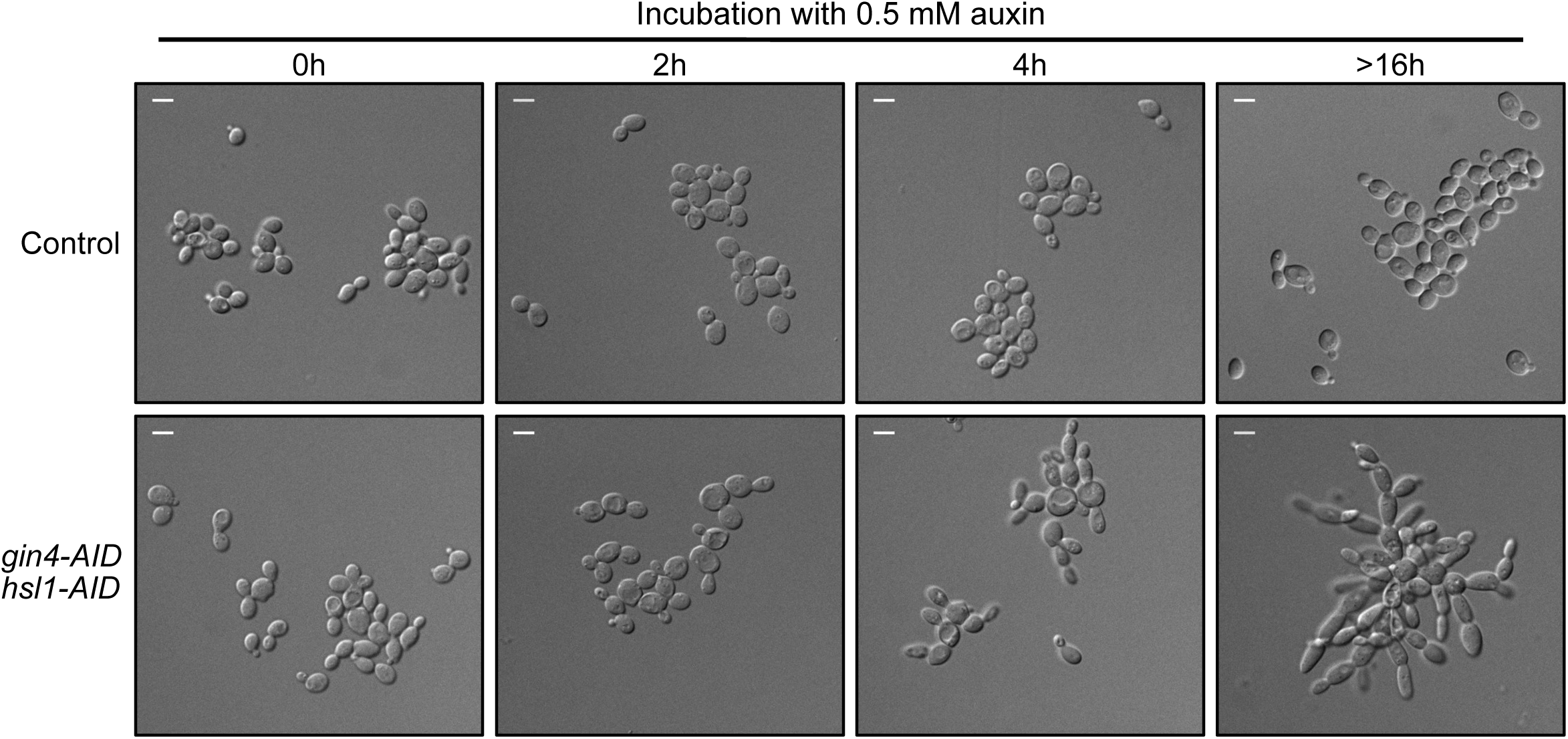
The severity of the *gin4-AID hsl1-AID* phenotype increases with time. Control cells and *gin4-AID hsl1-AID* cells were grown to log phase at room temperature in YPD medium and auxin was added to both strains. Both strains included 2 copies of the *TIR1* gene. DIC images of the cells were taken at the indicated times. Scale bar represents 5 µm.

**Figure S4:**
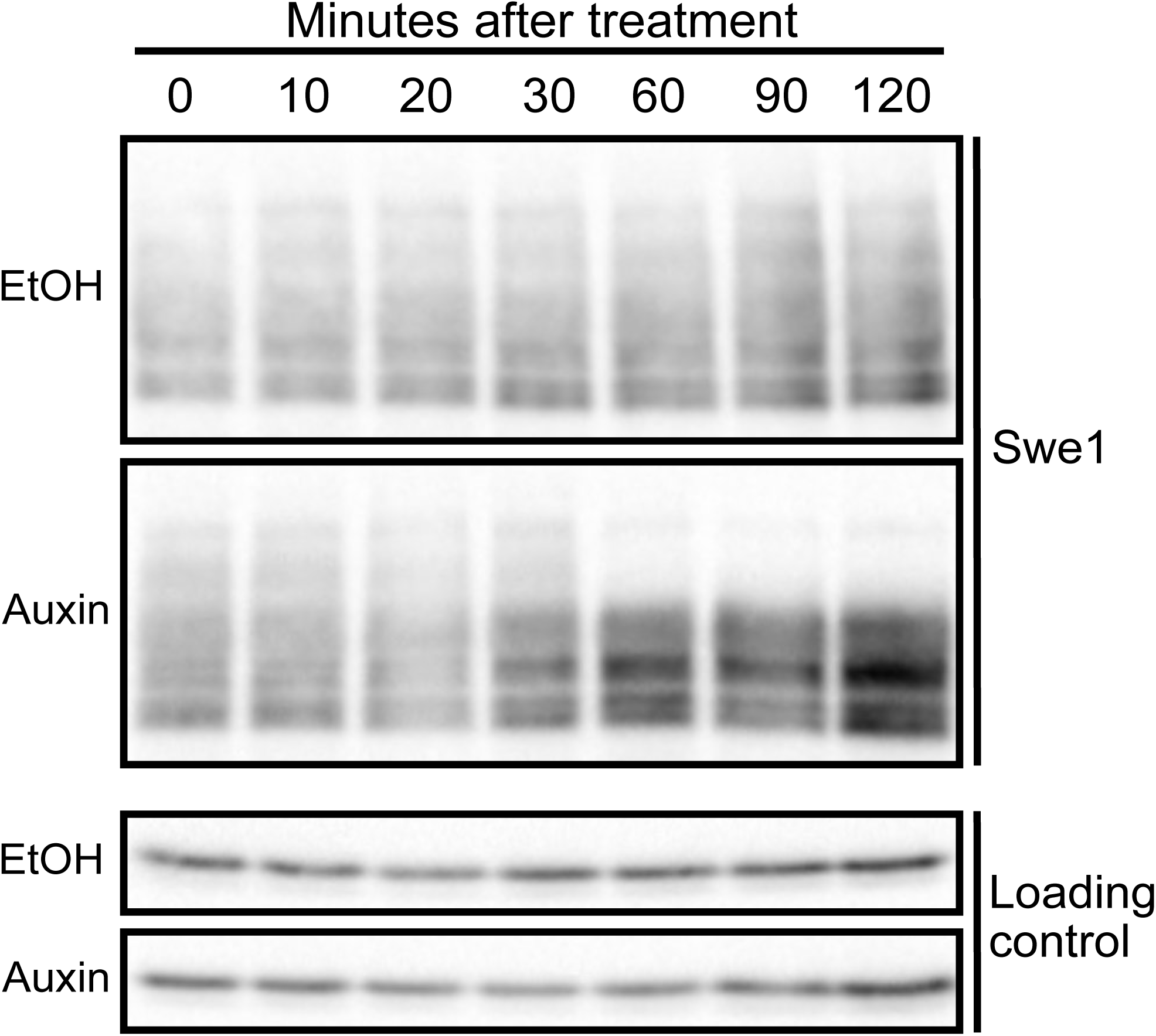
Swe1 is rapidly dephosphorylated in asynchronous cells upon inactivation of Gin4 and Hsl1. *gin4-AID hsl1-AID 2xTIR1* cells were grown to log phase in YPD and were then split into two aliquots. 0.5 mM auxin was added to one aliquot and an equivalent amount of the solvent for auxin was added to the other. Samples were taken at the indicated intervals and the behavior of Swe1 was analyzed by western blot. Anti-Nap1 was used as loading control.

**Figure S5:**
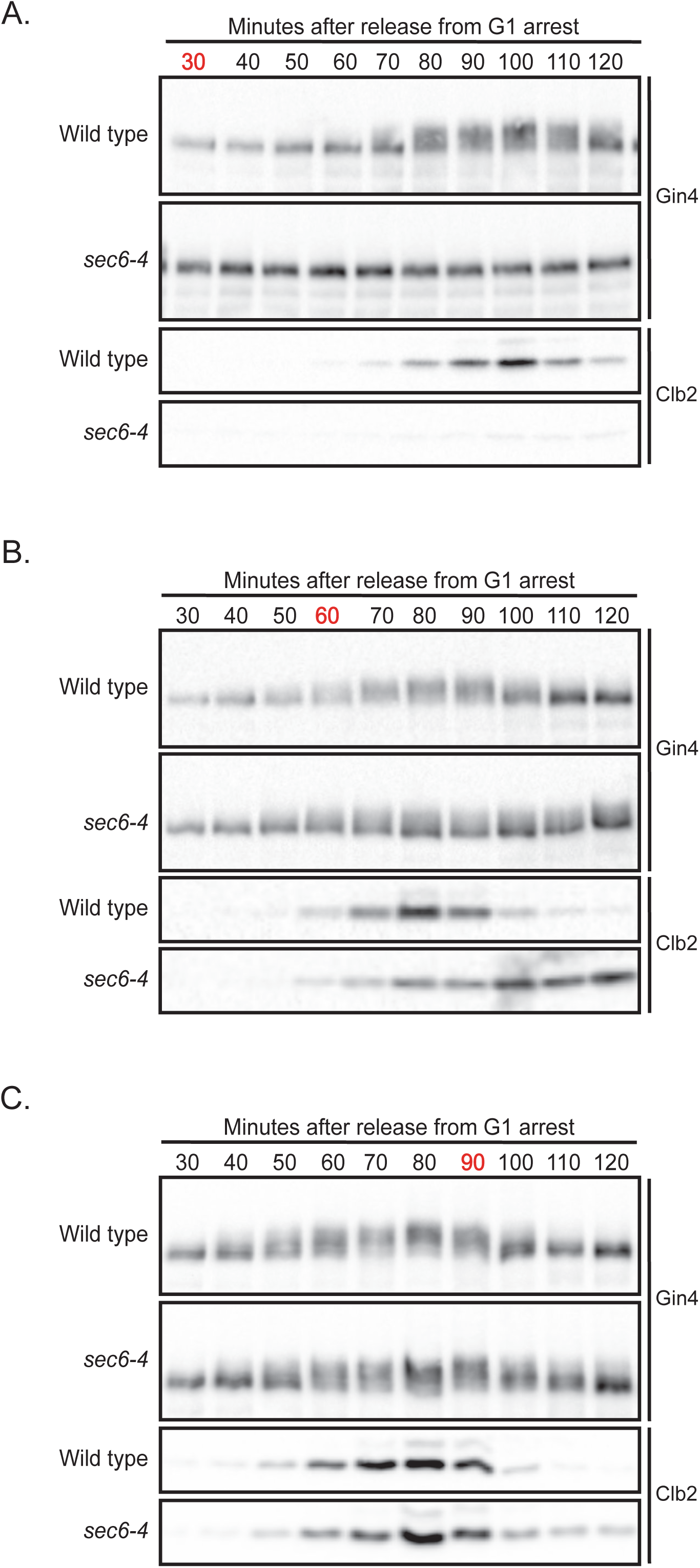
Gin4 phosphorylation is correlated with the extent of bud growth, rather than the rate of bud growth. Wild type and *sec6-4* cells were released from a G1 arrest in YPD at room temperature, incubated in a 25°C shaking water bath until a shift to the restrictive temperature (34°C) at 30 min **(A)**, 60 min **(B)** or 90 min **(C)** after release from arrest. Samples were taken at the indicated intervals and the behavior of Gin4 and Clb2 was analyzed by western blot. The timepoint labeled in red indicates the time at which cultures were shifted to 34°C.

**Figure S6:**
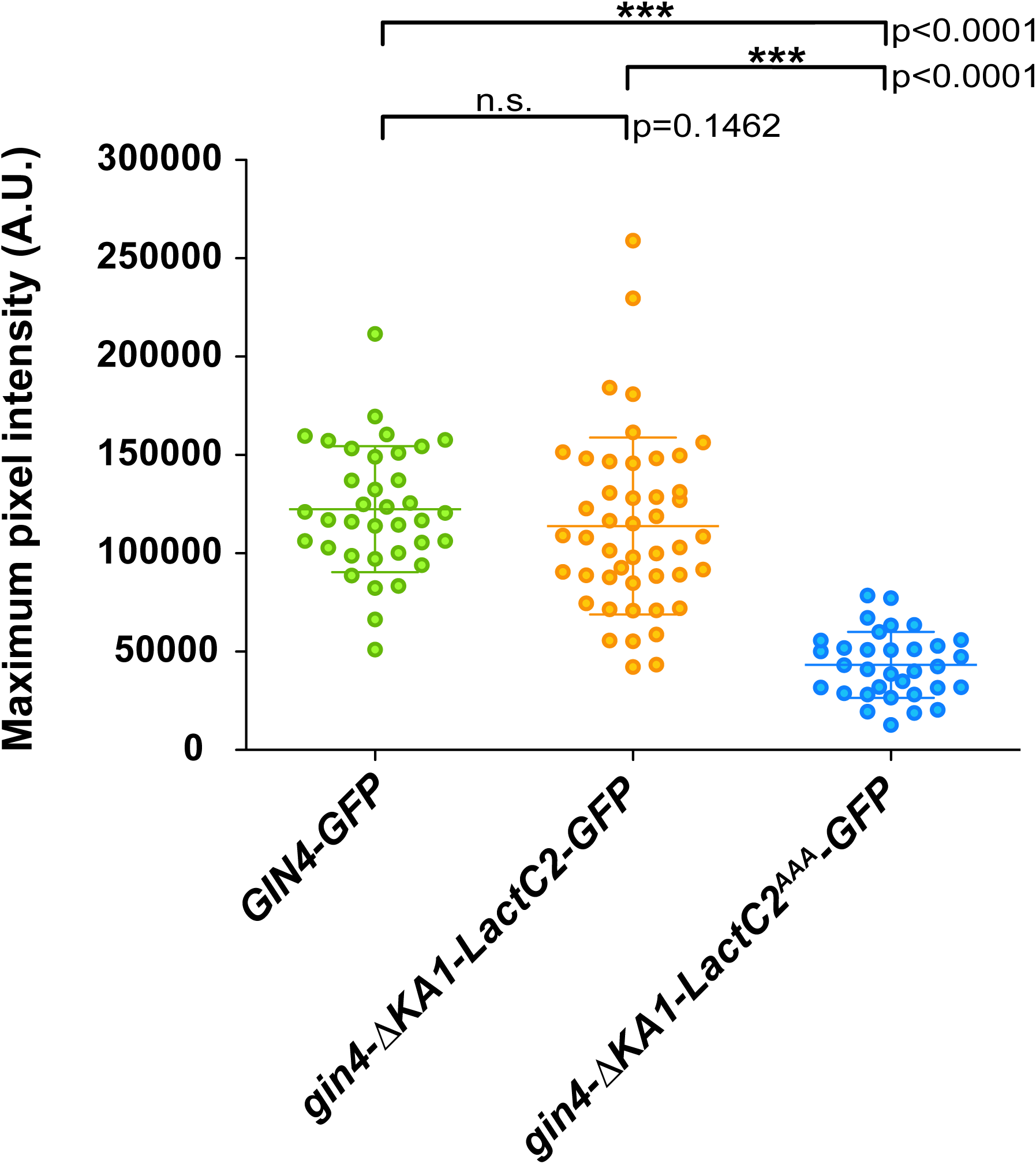
Quantification of Gin4-GFP localization at the bud neck. Cells of the indicated genotypes were analyzed to determine the maximum pixel intensity for GFP fluorescence at the bud neck. The Y-axis on the scatter plot shows the maximum pixel intensity in arbitrary units after subtracting background signal.

**Video 1: Time-lapse imaging of *2xTIR1* and *gin4-AID hsl1-AID* cells.** The two strains were mixed together prior to imaging as described in Materials and methods. The spindle pole bodies were differentially tagged in the two strains. The cells in top right corner show *2xTIR1* cells with mRuby2-tagged SPC42 and the cells in the bottom left corner show *gin4-AID hsl1-AID* with GFP-tagged SPC42. The *gin4-AID hsl1-AID* cells undergo a prolonged metaphase delay with polarized bud growth while the *2xTIR1* cells begin the next cell cycle. The cells were imaged using time-lapse confocal microscopy with image acquisition every 3 min. The movie was converted to AVI format using Fiji and shows the time-lapse at a speed of 5 frames per second (fps). Scale bar represents 5 µm.

**Video 2: *gin4-AID hsl1-AID* cells show spindle pole and cytokinesis defects.** A tile showing *2xTIR1* cells with mRuby2-tagged SPC42 and *gin4-AID hsl1-AID* cells with GFP-tagged SPC42 imaged together. Cells were imaged using time-lapse confocal microscopy with image acquisition every 3 min. The *gin4-AID hsl1-AID* cells exhibit defects in bud separation and results in the formation of cell chains. These defects are also accompanied by spindle defects (magenta arrows) in the second cell cycle. The movie was converted to AVI format using Fiji and shows the time-lapse at a speed of 5 fps.

